# Pathogenic and genetic diversity of *Sclerotium rolfsii,* the causal agent of Southern blight of common bean in Uganda

**DOI:** 10.1101/2025.08.15.670461

**Authors:** Samuel Erima, Moses Nyine, Mildred Ochwo Ssemakula, Geoffrey Tusiime, Emmanuel Amponsah Adjei, Eduard Ahkunov, Alina Akhunova, Settumba B Mukasa, Michael Hillary Otim, Thomas Lapang Odong, Allan Nkuboye, Agnes Candiru, Pamela Paparu

## Abstract

*Sclerotium rolfsi* Sacc. is a soil born fungus that causes southern blight to many crops in the tropical and subtropical regions. In 2018, Southern blight was reported to be the most prevalent bean root rot in Uganda. Earlier studies ascertained the morphological and pathogenic diversity of *S. rolfsii*, but limited understanding of its genetic diversity exists. Knowledge of *S. rolfsii* genetic diversity is a critical resource in the development of common bean varieties with durable resistance to the pathogen. A total of 188 *S. rolfsii* strains were collected from seven agro-ecological zones of Uganda in 2013, 2020 and 2021, and characterized for pathogenecity, growth rate, the number and colour of sclerotia produced and colony texture. The genetic diversity of the 188 strains was assessed using single nucleotide polymorphisms (SNPs) obtained from whole genome sequencing. The growth rate of the strains ranged between 1.1 to 3.6 cm/day while the number of sclerotia produced ranged between 0 to 543 per strain. The strains had fluffy, fibrous and compact colony texture. The sclerotia were either brown or dark brown. The analyzed strains were pathogenic on common bean and caused a disease severity indices between 10.1% and 93.3%. Average polymorphic information content across all chromosomes was 0.322 with a minimum of 0.145 and maximum of 0.525. Average observed heterozygosity across all the chromosomes was 0.782. Population structure analysis identified five genetically distinct groups. The results of analysis of molecular variance revealed that 0.39% of the variation was between population while 11.98% was between samples within population and 87.62% between samples across populations. The average genetic distances between the isolates from different agro-ecological zones was 0.038 while the average similarity coefficient was 0.968 There were significant differences in the morphological traits and pathogenicity across the different genetic clusters. This high levels of genetic diversity within strains needs to be taken into consideration while selecting strains for screening breeding lines.

**Key word:** Pathogenicity, genetic diversity, Characterisation, Southern blight, common bean

## Introduction

The common bean (*Phaseolus vulgaris* L.) is the third most important legume in the world after soybean (*Glycine max* L.) and peanuts (*Arachis hypogaea* L.) (Miklas *et al.,* 2006; Broughton *et al.,* 2003). In Uganda, it is the third most important annual crop after maize (*Zea mays* L) and Cassava (*Manihot Esculentus* C) (UBOS, 2022). In 2019, common bean production declined to 438,000 tons from 1,008,410 tons in 2016 yet area planted had increased to 800,000 hectares from 670,737 hectares with average yield of 0.6 tons per hectare (UBOS, 2022; FAOSTAT, 2018). Uganda Bureau of Statistics (UBOS) attributed this decline to reduced productivity. Factors known to negatively influence common bean productivity include; poor soil fertility, pests (bean stem maggot, aphids, bean leaf beetles, etc.) and diseases (root rots, angular leaf spot and anthracnose, etc.), poor agronomic practices, weed competition, moisture stress and lack of quality seed (Gatsby, 2014; Sinclair and Vadez, 2012).

*Sclerotium rolfsii* Sacc. is a soil borne pathogen that primarily attacks the root and stem, but can also attack the pods, petioles and leaves when these plant parts are in touch with the soil (Kator *et al.,* 2015). The species was first described in 1911 by Italian mycologist Andrea Saccarda but it was not until 1982 when it was first identified as the causal agent of Southern blight in tomatoes (*Solanum lycopersicum* L.) (Rolfs, 1982). *Sclerotium rolfsii* is the anamorphic stage (asexual state) of the pathogen while *Athelia rolfsii* is the teleomorph state (sexual state) and is rarely observed. The pathogen is classified as follows; Kingdom-Fungi, Phylum-Basidiomycota, Subphylum-Agaricomycotina, Class-Agaricomycetes, Order-Atheliales, Family*-Atheliaceae* and genus *Athelia* (Kator, 2015). *Sclerotium rolfsii* is commonly found in the tropics, sub tropics and warm temperate regions especially in the United States, Central and South America, West Indies, Southern Europe, India, Japan, the Phillipines and Hawai. The pathogen rarely occurs when temperatures drop below 0°C (Kator *et al.,* 2015).

Paparu *et al.,* (2018) reported Southern blight, as the most prevalent bean root rot in Uganda besides Fusarium, Pythium and Rhizoctonia root rot diseases. The pathogen causes both pre- and post-emergence dumping off, with the former mostly observed in highly susceptible varieties and for highly virulent strains. For post-emergence dumping off, wilting can occur as early as one week after planting (Paparu *et al.,* 2020). There is general wilting of the plant characterized by dry shoot and plant death. Under conditions of high humidity, abundant white mycelia are usually visible on the collar and root of infected plants. Brown sclerotia are usually visible at the base of the plant at advanced stages of wilting (Ferry and Dukes, 2002). Yield losses due to Southern blight in the field range between 55.5 and 64.9% (Alam *et al.,* 2018), yet in Uganda there is no common bean variety known to be resistant to the pathogen (Paparu *et al*., 2018). Use of host plant resistance is the most cost effective and sustainable strategy for the management of root rots by the smallholder farmers who are the major bean growers in sub saharan Africa (Mabaya *et al*., 2020; CIAT, 2003). And knowledge of pathogen diversity is key in efforts to develop durable resistance in crops.

Pathogen diversity can be determined using morphological/cultural, pathogenic and genetic studies. Morphological/cultural divsersity studies are often cheap, but they may not distinguish strains that produce similar morphological structures and they require experienced mycologistis to conduct them (Trabelsi et al., 2017). Pathogenic diversity studies aid in determing the host range of the pathogen and the selection of strains which can be used for screening the germplasm. (Le et al., 2012)

Genetic diversity of *S. rolfsii* has been studied using PCR based molecular markers including Internal Transcribed Spacers (ITS) region of the ribosomal RNA (Amaradasa *et al*., 2020), Translation Elongation Factor 1 alpha gene region (TEF 1 α) (Chen *et al*., 2020), Randomly Amplified Polymorphic DNA (RAPD) (Kulkami and Hedge, 2019) and Amplified Fragment Length Polymorphism (AFLP) (Cilliers *et al*., 2000). However, these amplicon sequencing (AmpliSeq) approaches require up front variant discovery to design and synthesize the primer sets (Adhikari *et al*., 2022). In the last decade, next generation sequencing (NGS) has been utilized (Yan *et al*., 2021; Iquabel *et al*., 2017) with approaches such as genotyping by sequencing (GBS) and restriction site associated DNA sequencing (RAD Seq) used for diversity studies. Though the RAD Seq and GBS do not require prior sequence information, however, they are characterized by reduced representation of the genome (Kumar *et al*., 2021). Due to advances in the NGS technologies and reduction in the cost of sequencing, skim sequencing (SkimSeq) of the whole genome at low coverage can be used to identify large number of polymorphic markers cost effectively (Adhikari *et al*., 2022; Kumar *et al*., 2021). Availability of abundant single nucleotide polymorphism (SNP) markers facilitates diversity studies in organisms (Wang *et al*. 2023). However, to apply next generation sequencing, high quality DNA is needed. Obtaining high quality DNA from *S. rolfsii* is reported to be challenging due to the presence of extracellular polysaccharides in their cell walls (Male *et al*., 2018). However, some modifications in the DNA extraction methods are reported to effectively remove these polysaccharides (Male *et al*., 2018; Pérez *et al.,* 2013; Jeeva *et al.,* 2008).

Although, Paparu *et al.,* (2018 and 2020) studied the morphological and pathogenic diversity of *S. rolfsii* strains collected before *2020*, the genetic diversity of these strains was never characterized. Therefore, this study intended to 1) Determine morphological diversity of strains not previously studied, 2) Determine the pathogenicity of all *S. rolfsi* strains following long periods of storage, and 3) Conduct genetic diversity and population structure analyses of the *S. rolfsii* strains using whole-genome re-sequencing data. The genetic differentiation among the sub-populations of *S. rolfsii* was compared with agro-ecology of strains’ origin, growth rate, and numberof sclerotia produced to assess their potential role in pathogen adaptation and virulence on comon bean The generation of these information can facilitate pathogen suveylance and selection of isolates to screen germplasm for breeding disease resistant varieties.

## Materials and methods

### Morphological diversity of selected *S. rolfsii* strains

Eighty-four (84) *S. rolfsii* strains isolated in the years 2020 and 2021, and not previously characterized morphologically were used in this study. The pathogen was isolated according procedure to the procedure by Paparu et al. (2020). Morphological diversity was studied by assessing the growth rate, colony characteristics and the number of Sclerotia produced by each strain. Isolates that had been stored on filter papers were initiated on PDA (Esvee Biologicals, Mumbai, India). The Isolates were purified by hyphal tipping according to procedure by Paparu et al. (2020). We used Potato dextrose agar (PDA) with 0.3 g of rifamycin per liter to conduct morphological characterization according to the procedure by Paparu et al. (2020). Inoculation was performed by touching the surface of the fungal culture on a petri dish with a needle and then transferring the attached mycelia to the centre of the Petri dish. Each fungal strain was inoculated in triplicate onto 8-cm diameter Petri dishes and incubated in the dark at 25 °C. Colony diameter was measured daily, beginning 2 days post-inoculation, and continued until the mycelium fully covered the plate surface, which occurred by day 4 for most strains. Following growth assessment, colony color and texture were documented. Cultures were maintained for 28 days, after which the number of sclerotia were recorded following the protocol described by Mullen (2001).

### Pathogenic diversity of *S. rolfsii* strains

#### The *S. rolfsii* strains

The pathogenicity of 188 *S. rolfsii* strains including 104 strains previously studied by Paparu *et al*. (2020), and the 84 strains collected in 2020 and 2021 was investigated in this study. Pure cultures of these strains preserved on filter paper and stored in Eppendorf tubes at 4°C were tested for pathogenicity for the first time following long period in storage. The strains were collected from several agro-ecological zones including the Eastern Highlands (EH), Lake Victoria Crescent and Mbale farmlands (LVC), Northern Mixed Farming System (NMFS), South Western Highlands (SWH), Teso Farming Zone (TFZ), Western Mixed Farming System (WMFS) and West Nile Mixed Farming Zone (WNMFS). Three strains out of the 188 were isolated from Bukoba in Tanzania.

#### Preparation of *S. rolfsii* inoculum

The analyses of *S. rolfsii* pathogenicity were conducted according to the procedure by Paparu *et al*. (2020). A total of 188 strains were screened against five common bean lines: MLB49-89A (Fusarium root rot resistant check), RWR719 (Pythium root rot resistant check), KWP9 (Southern blight tolerant check), NABE 14 (released variety, tolerant to Fusarium root rot Geosttch *et al*. (2016) and CAL96, also known as K132 (susceptible check for common bean diseases).

In brief,50g of millet were mixed with 50ml of water in an autoclave bag and sterilized by autoclaving at 121 °C for 1 hour. Five 1-cm² agar plugs were excised from 14-day-old PDA cultures (39 g/L) grown in 9 cm Petri dishes. The plugs were aseptically mixed with sterilized millet in a laminar flow cabinet, sealed in bags, and incubated at 25 °C for 21 days.

#### Soil inoculation and planting of common bean lines

Twenty kilograms (20 kg) of steam-sterilised loam soil and sand mixed in the 2:1 ratio was placed in the 70 cm × 35 cm × 10 cm wooden trays lined with a polythene sheet after disinfecting with 75% ethanol. The soil was then inoculated with 10 g of the millet carrying fungal mycelia and sclerotia. Each strain was inoculated in a single tray and replicated once. Trays were arranged in a Completely Randomized Design (CRD) in the screen house. In each tray, 16 seeds of each line were planted in two rows. Common bean lines were randomized in each tray in a split plot design. Sixteen seeds of each of the test lines planted in an un-inoculated tray were used as controls. The planting trays were protected from direct rainfall, but watered twice daily until the experiments were terminated.

#### Assessment of seed germination, Southern blight incidence and severity

At 14 days after planting, germination data were collected. The seeds that did not germinate were examined to assess for pre-emergence dumping off (characterized by rotting and fungal growth on seed) and non-viable (intact seed with no sign of rotting or fungal growth). Plants that germinated were assigned germination score 1 while those that did not germinate were assigned score 0. Preliminary data on incidence and severity was also collected at this point where plants that died of pre-emergence dumping off were assigned score 1 for incidence and severity score 6 for pre-emergence dumping off. Final Southern blight severity was assessed 28 days after planting using a scale developed by Le *et al.,* (2012) and modified by Paparu *et al*. (2020) by adding a score 6 to represent pre-emergence dumping off (Figure 1).

**Figure 1.**
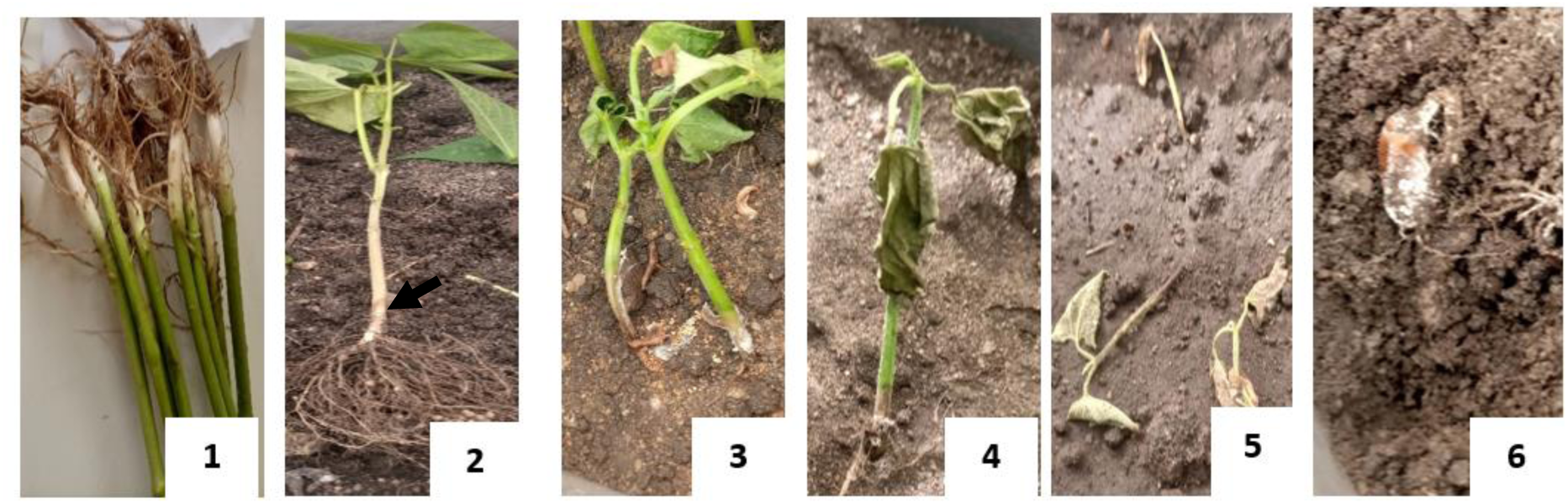
Southern blight disease severity index scale (1-6): 1 = No symptoms, 2 = grey water-soaked lesions on stem above soil but no visible fungal growth, 3 = visible fungal growth at the base of the stem near the soil line, characterized by silky white mycelia or sclerotia that gradually darken, 4 = partial wilting where young leaves begin to wilt and stems begin to shrivel, 5 = complete wilting, desiccation and browning of leaves and stem collapse and death of plant, and 6 = pre-emergence damping off, complete seed death characterized by rotting and fungal growth.

#### Genetic diversity of *S. rolfsii* strains

*Sclerotium rolfsii* produces extracellular polysaccharides which co-precipitate with DNA adversely affecting DNA recovery (Schmid *et al*, 2013). Male *et al*. (2018) also reported that DNA extraction kits do not work well for *S. rolfsii*. Hence a modification of the protocol by Joint Research Centre (JRC) European Commission (2024) was adopted where dry mycelia was used and precipitated on ice using absolute isopropanol and 2.5M Sodium acetate.

A DNA extraction buffer (2 M NaCl, 0.2 M EDTA, 0.2 M Tris HCL (PH8) and 1% SDS) was prepared and autoclaved. Mycelia from two-week old culture were scrapped into sterile Petri dishes and oven-dried overnight at 35 °C. About 0.02 g of the dry mycelia was put in 2-ml Eppendorf tubes containing beads and ground at 1400 revolutions per minute for 3 minutes in a Geno grinder (1600 MiniG, Cole-Parmer, Chicago, USA). Then 800 µl of DNA extraction buffer preheated to 65 °C was added to the ground mycelia and ground for another 2 minutes in the Geno grinder. The tubes were then incubated in a water bath at 65 °C for one hour, followed by centrifugation at 12,000 revolutions per minute (rpm) for 10 minutes. The supernatant (500 µl) was transferred into sterile 2-ml Eppendorf tube. An equal volume of chloroform and isoamyl alcohol (24:1) was added, mixed for 2 minutes and centrifuged at 12,000 rpm for 10 minutes to separate the DNA from the cellular residues. If a clear supernatant was not obtained, this step was repeated. The supernatant (400 µl) was transferred into a 1.5-ml sterile Eppendorf tube, mixed with An equal volume of ice-cold isopropanol stored at -20 °C by inverting a tube 5 to 10 times. Then 40 µl of 2.5 M sodium acetate was added to each tube and inverted once.

The solution was incubated at -20 °C for two hours to precipitate DNA. Samples were centrifuged at 12000 rpm for 10 min to pellet the DNA and the supernatant was discarded. The pellets were washed twice with 500 µl of 75% ethanol and left to air dry at room temperature for 1 hour. After this the DNA was dissolved in 100 µl of 1x TE buffer (10 mM Tris HCL (pH 8), 1 mM EDTA). DNA was quantified using a NanoDrop (ND-1000) (Thermo Fisher, Waltham, Massachusetts, USA). The integrity and quality of DNA was assessed using electrophoresis with 1% w/v agarose gel. The extracted DNA was shipped to Kansas State University where further assessment of the DNA quality and integrity was performed using a TapeStation 4200 (Avantor, Radnor Pennsylvania, USA).

Libraries were prepared using Nextera DNA library preparation protocol (<u>Nextera DNA</u> <u>Library Prep Protocol Guide (1000000006836) (https://www.illumina.com) with reduced reaction volume</u>. (Adhikari et al., 2022; Jordan et al., 2022) The libraries were subjected to size selection using the Pippin Prep system (Sage Scientific, Beverly, MA, USA) to enrich for 400–600 bp fragments. Barcoded libraries were pooled and sequenced on NovaSeq 6000 to generate ∼16.5x coverage of 2 x 150 bp paired-end reads per sample.

### Data analysis

#### Analysis of germination, incidence and severity data

Percentage germination for the 16 plants of each common bean variety was calculated in Microsoft excel. The severity of Southern blight was calculated according to the formula by Chiang *et al.,* (2017).

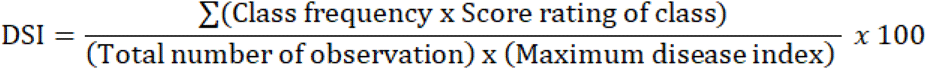

Analysis of variance (ANOVA) was used to determine the effect of strains on pathogenicity on common bean varieties. The following ANOVA model was fit to the Southern blight severity score data: Y_ij_ =µ+g_i_+e_ij_, where, Y_ij_ is the observed severity of the i^th^ genotype in j^th^ experiment, μ is the overall population mean, g_i_ is the genetic effect of the i^th^ variety, and e_ij_ is the effect of experimental error. Tukey’ honestly significant difference (HSD) was used to compare group means. The Pearson’s chi square test (X^2^) was used to test for association between pathogenicity and geographical origin of the strains.

#### Analysis of S. rolfsii sequence data

Sequence reads were demultiplexed using a custom python script to separate individual sample reads based on the index barcodes. The adaptor trimming and read quality filtering was performed using Trimmomatic v0.32 software (Bolger *et al*., 2014). The high-quality reads were aligned to the indexed reference genome of *Agroathelia rolfsii* ASM1834391v1 (Yan et al., 2021) using Burrows-Wheeler Alignment Tool (bwa-mem) (Li *et al*., 2009). The samtools software was used to convert alignments from sam to bam formats and Picard tools (Broad Institute, 2019) were used to add read groups and sample IDs. Further processing was done using the genome analysis tool kit (GATK) HaplotypeCaller to generate the genome variant call format (gvcf) files for each sample. The gvcf files were combined using GenomicsDBImport for variant calling using Genotype GVCFs from GATK (Van der Auwera, G. A., & O’Connor, 2020). The resulting variant call file was pre-filtered for minor allele frequency (>0.05), genotype missingness (<15%) and Insertions and deletions (INDELs) using vcftools v0.1.13. Additional filtering was done in Tassel (v5.2.52) at 0.05 minor allele frequency to retain SNPs that were used in the downstream analyses. Data was then imputed using K nearest neighbor genotype imputation method (Bradbury et al., 2007) and 51,447 SNPs were used for downstream analysis. The SNP diversity was summarized calculating the distribution of minor allele frequency (MAF) (Bradbury et al., 2007) and polymorphic information content and heterozygosity (PIC) (Liu et al., 2005). Sparse non-negative matrix factorization (SNMF) was used to create genetic clusters of the isolates for different values of K ranging from K = 1 to K = 10 to find the optimum number of clusters that can explain the genetic variation (Frichot et al., 2014). A cross-entropy plot was generated to identify the minimum number of clusters that can explain the variation between clusters. Principal component analysis (PCA) and discriminate analysis of principal component (DAPC) were also used to determine the population structure of the strains. The optimum number of genetic clusters was further verified in structure software v 2.3.4 according to procedure by Evanno et al., (2005).

Analysis of molecular variance (AMOVA) was conducted to assess the variation among the strains based on geographical origin and genetic clusters using “poppr” and “adegenet” package in R software (Kamvar et al., 2014; R core team, 2018). Genetic variation was portioned into variation within and between agro-ecological zones and genetic clusters. A phylogenetic tree was generated using Euclidean distance matrix in power marker v3.25 (Liu et al., 2005) and visualized using fig tree. “poppr” and “adegenet” package in R software were used to compute genetic distance and similarity coefficients between isolates from the different agroecological zones. Association between the clusters, geographical origin, pathogenicity, growth rate and number of sclerotia were tested using Chi square(X).

## Results

### Morphological diversity of *S. rolfsii* strains

Data were collected on the growth rate and number of sclerotia of the 84 strains sampled in 2020 and 2021. The growth rate of the strains ranged from 1.1 to 3.6 cm/day while the number of sclerotia produced ranged from 0 to 543. The number of sclerotia produced was categorized as none, low, medium and high as follows; 0 = none, 1 to 49 = low, 50 to 99 = Medium and > 100 = high. A total of 12, 21, 20 and 31 strains were in the groups of high, medium, low and none, respectively. The strains with cottony colony texture (Figure 3) were classified as fibrous (12 strains), compact (2 strains) and fluffy (70 strains). The sclerotia were brown for 40 strains and dark brown for 13 strains, while 31 strains did not form sclerotia by day 28 post inoculation. Significant differences were observed in the number of sclerotia produced by isolates sampled from different agroecologal zones (F = 5.6, P = 0.001). Following Tukeys, HSD test, the average number of sclerotia produced by strains from LVC, NMFS, WNMFS and Tanzania were similar while that of the rest of the agro-ecological zones differed (Figure 2A).

**Figure 2:**
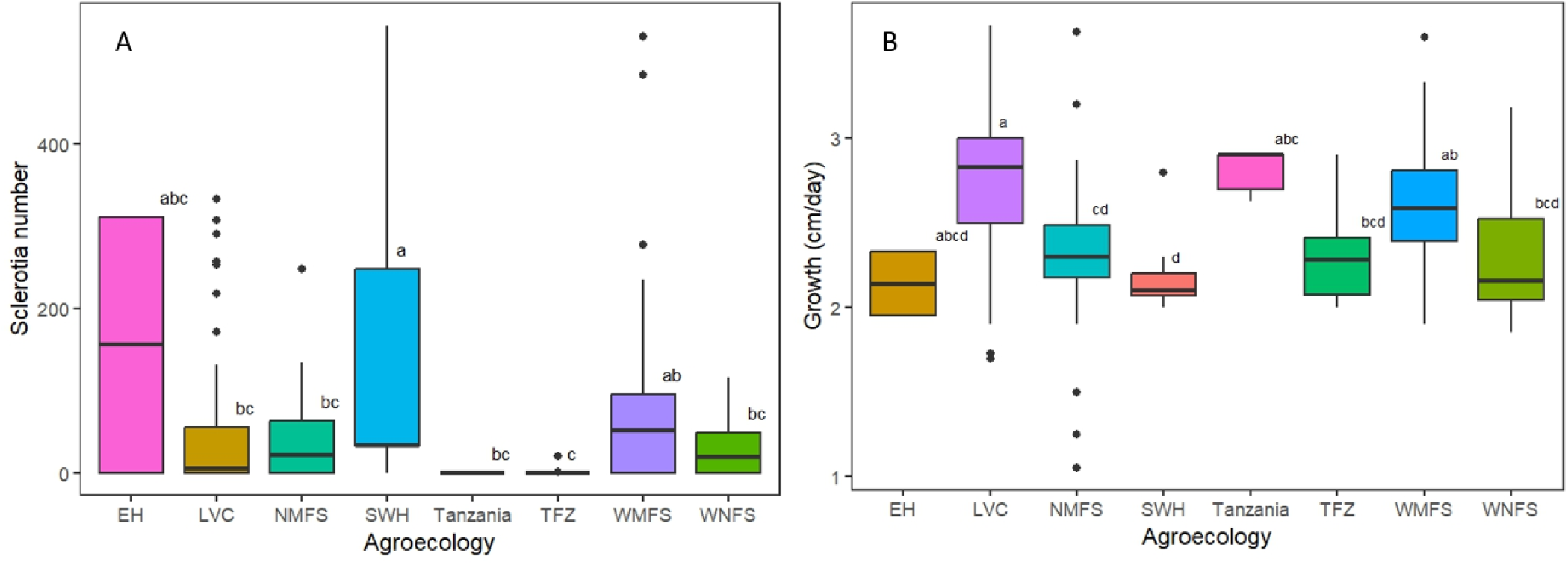
Comparison of the number of sclerotia and growth rates among the isolates sampled from distinct geographic locations. A. The average number of sclerotia. B. The growth rate of different S. rolfsii strains and their agroecologies of origin

**Figure 3:**
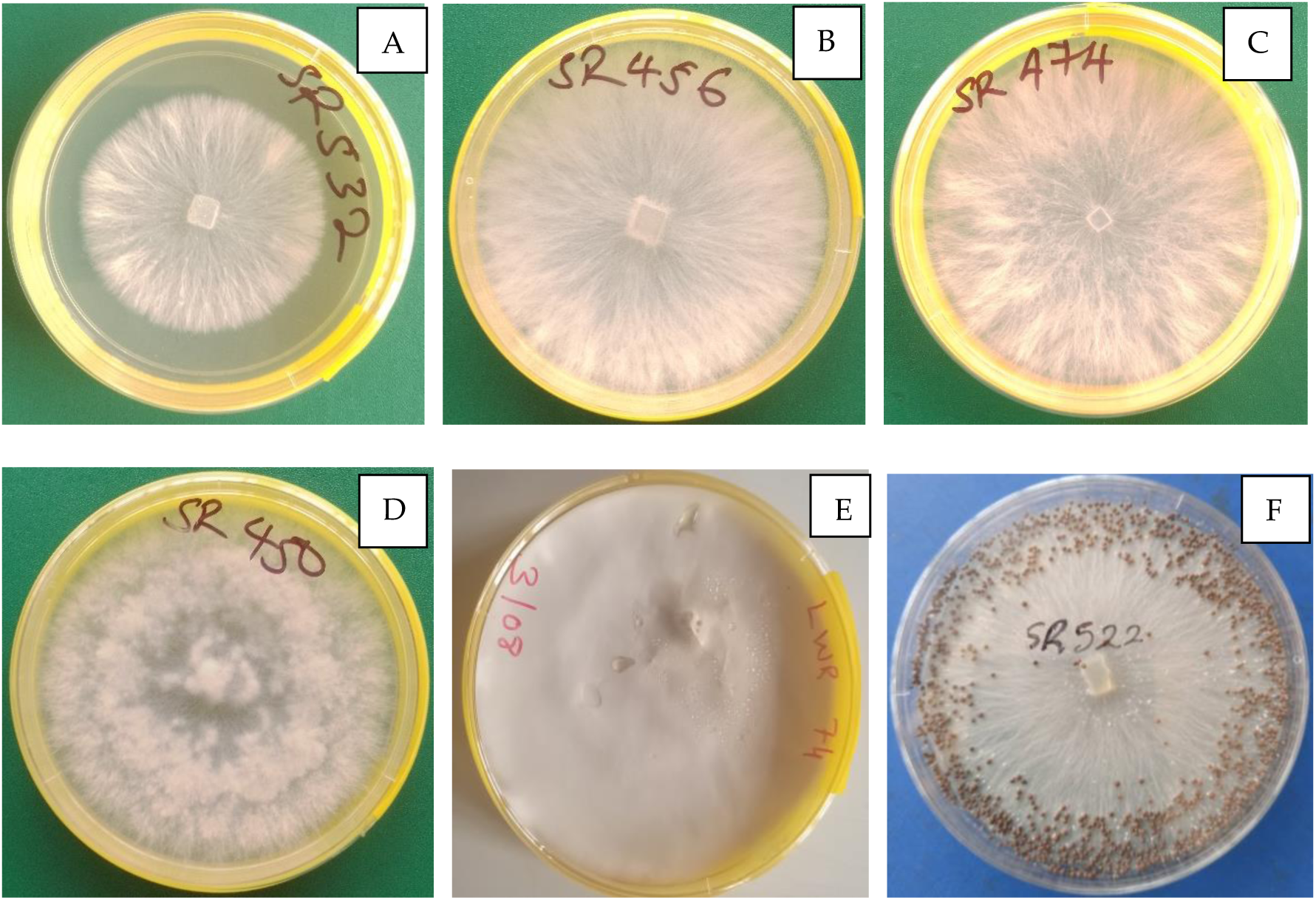
Colony morphology of *Sclerotium rolfsii* strains following culture on potato dextrose agar (PDA). A -C. Colonies produced by strains (SR532, SR456 and SR474) with the fibrous texture. D. Colony produced by strain SR450 with the fluffy texture. E. Colony produced by strain LWR74 with compact texture. F. Colony produced by strain SR522 showing numerous sclerotia.

Significant differences were also observed in the average growth rate of the strains from distinct sampling locations (F = 12, P = 0.0001) (Figure 2B). The strains from WNMFS and TFZ had similar average growth rates, while the strains from the rest of the agro-ecological zones differed in the growth rates (Figure 2B). The strains from Bukoba in Tanzania, had the highest average growth rate of 2.81±0.054 cm/day while those from EH had the lowest growth rate of 2.14±0.058 cm/day. Meanwhile, strains from EH had the highest average sclerotia number of 156 and the ones from TFZ had the lowest average sclerotia number of 2. The strains from Tanzania did not produce any sclerotia in the 28 days the experiment was ran. Additional data on the rest of the strains (104 collected in 2013) was obtained from Paparu et al. (2020) and summarized in table S1.

### Pathogenic diversity of *S. rolfsii*

#### Virulence of S. rolfsii strains

Pathogenicity tests were conducted on 188 strains. Of these, 104, 12 and 72 were collected in 2013, 2020 and 2021, respectively (Table S2). The Disease severity index (DSI) was evaluated for each strain grouped into 5 experimental groups, with experiment 1 including 58 isolates, experiment 2 including 45 isolates, experiment 3 including 26 isolates, experiment 4 including 28 isolates and experiment 5 including 31 isolates. DSI was significantly different among *S. rolfsii* strains in experiment 1 (F= 2.93, *P* = 0.0001), experiment 2 (F = 2.67, *P* = 0.0009) and experiment 4 (F = 2.186, *P* = 0.028). However, there was no significant difference in the DSI of the strains in experiment 3 (F = 1.27, P = 0.26) and experiment 5 (F = 1.43, P = 0.38). When the means were compared using the Tukey’s HSD, in experiment 1, strains SR282, SR270, SR285, SR208, SR210, SR205, SR308, SR348, SR200 and SR264 had significantly different DSI from the rest of the strains. In experiment 2, strains SR406, SR52, SR59, SR56, SR410, SR48, SR37, SR35, SR45 and SR9 had significantly different DSI from the rest of the strains. The Tukeys’s HSD did not show significant differences among the mean DSI of strains in experiment 4.

Overall, the most virulent strain was SR282 from West Nile Mixed Farming System (WNMFS) with an average DSI of 93.3%. The least virulent strains were SR475 and SR489 from Sironko district in the Lake Victoria Crescent and Mbale Farmlands Agorecology (LVC) with DSI of 10.1%. Out of the 10 most pathogenic strains, 8 were from 2013 collection while only two were from 2020/2021 collection (Table 1 and Table S1). The average DSI of 2013, 2020 and 2023 were significantly different from each other (F = 17.2, P = 0.001) (Figure 4B). The average DSI of 2013, 2020 and 2021 strains across all the experiments were 59.5%, 46.3% and 48.8%, respectively (Figure 5). Out of the 10 least virulent strains, 7 were from the 2020/2021 collection while only three were from 2013 collection (Table 2 and Table S2). The average DSI of strains for the different agro-ecological zones ranged between 46.6% for Teso Farming System (TFZ) to 65.1% for South Western Highlands. The average DSI of isolates from Bukoba in Tanzania was 76.8%, which was the highest among all the geographical locations (Figure 4A and Table S3). Chi square revealed no association between year of storage and virulence (X = 329, P = 0.351).

**Figure 4:**
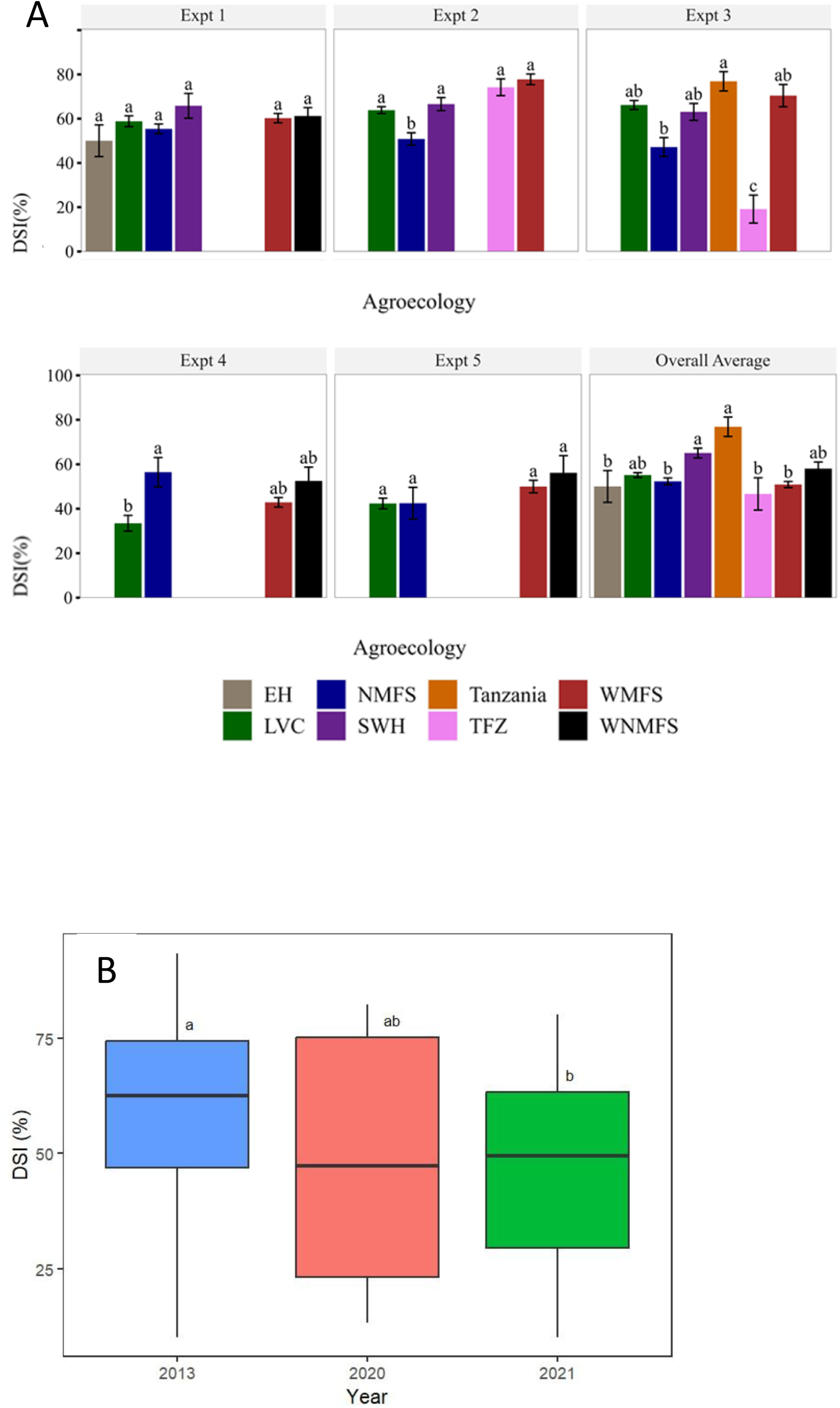
A Disease severity index caused by different *S. rolfsii* strains from eight common bean agro-ecological zones across five experiments. EH = Eastern Highlands, LVC = Lake Victoria Crescent and Mbale Farmland, NMFS = Northern Mixed Farming System, SWH = South Western Highlands, TFZ = Teso Farming Zone, WMFS = Western Mixed Farming System and WNMFS = West Nile Mixed Farming System. B-Disease severity index and the year of collection of the strains

**Figure 5:**
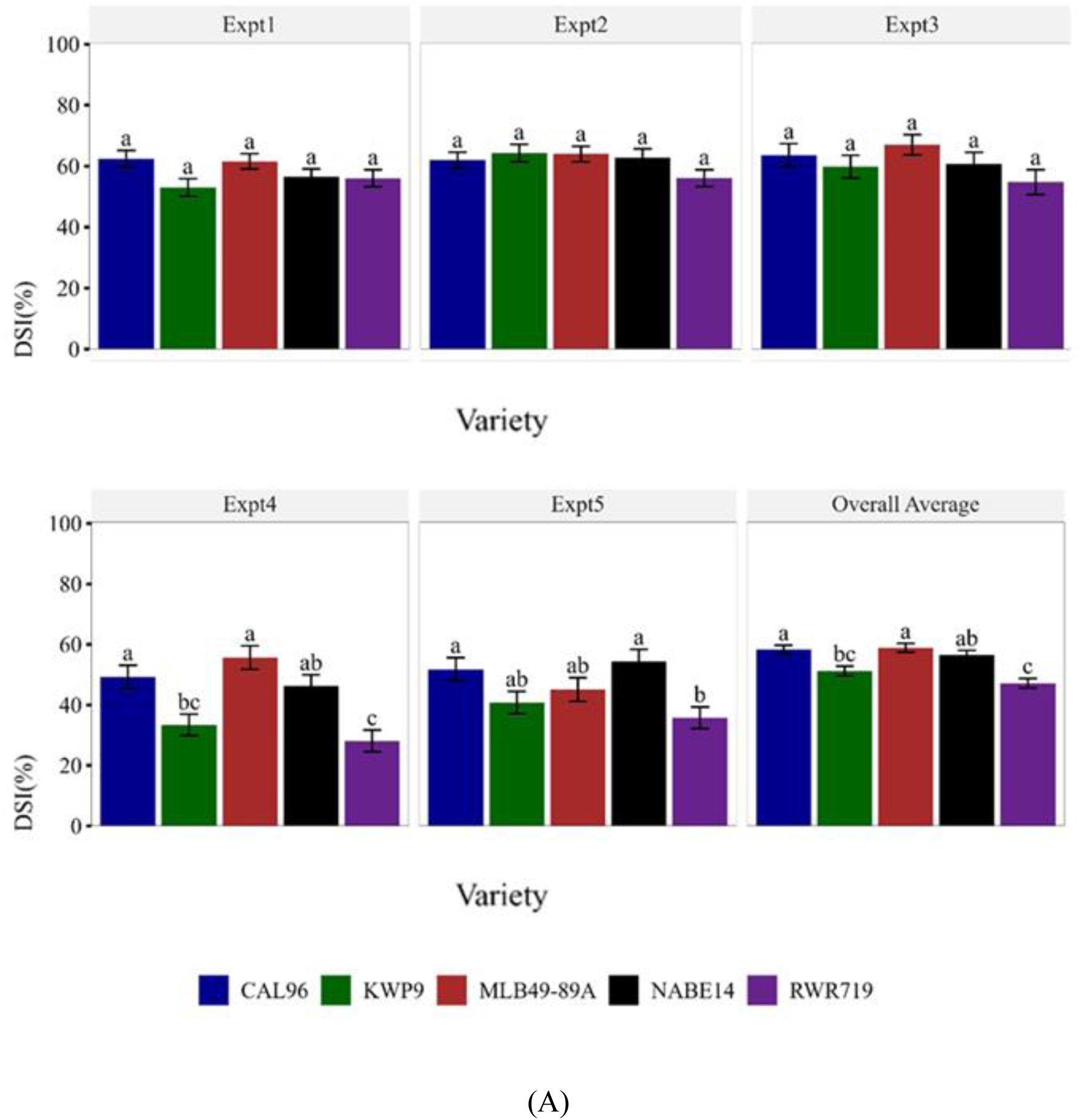
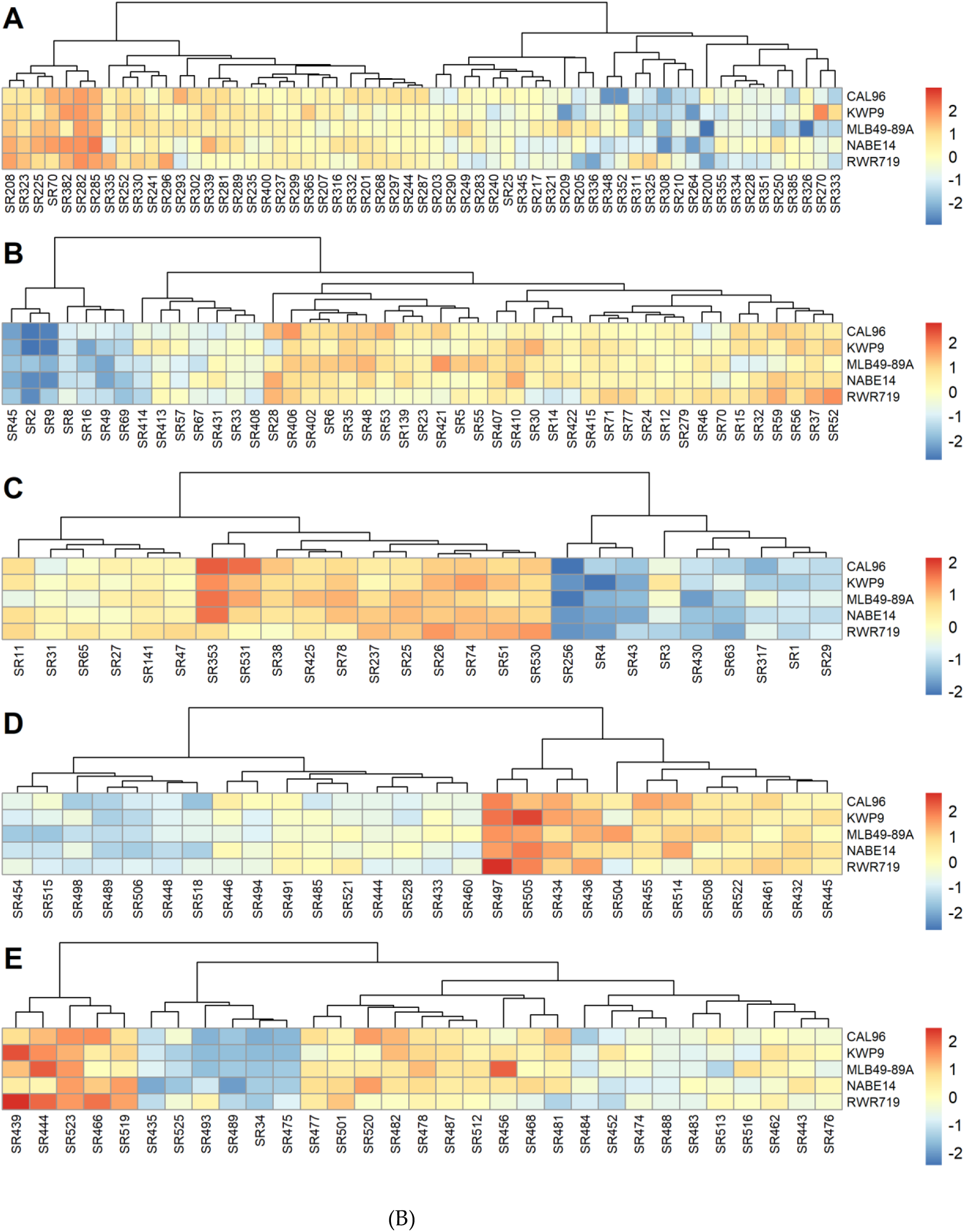
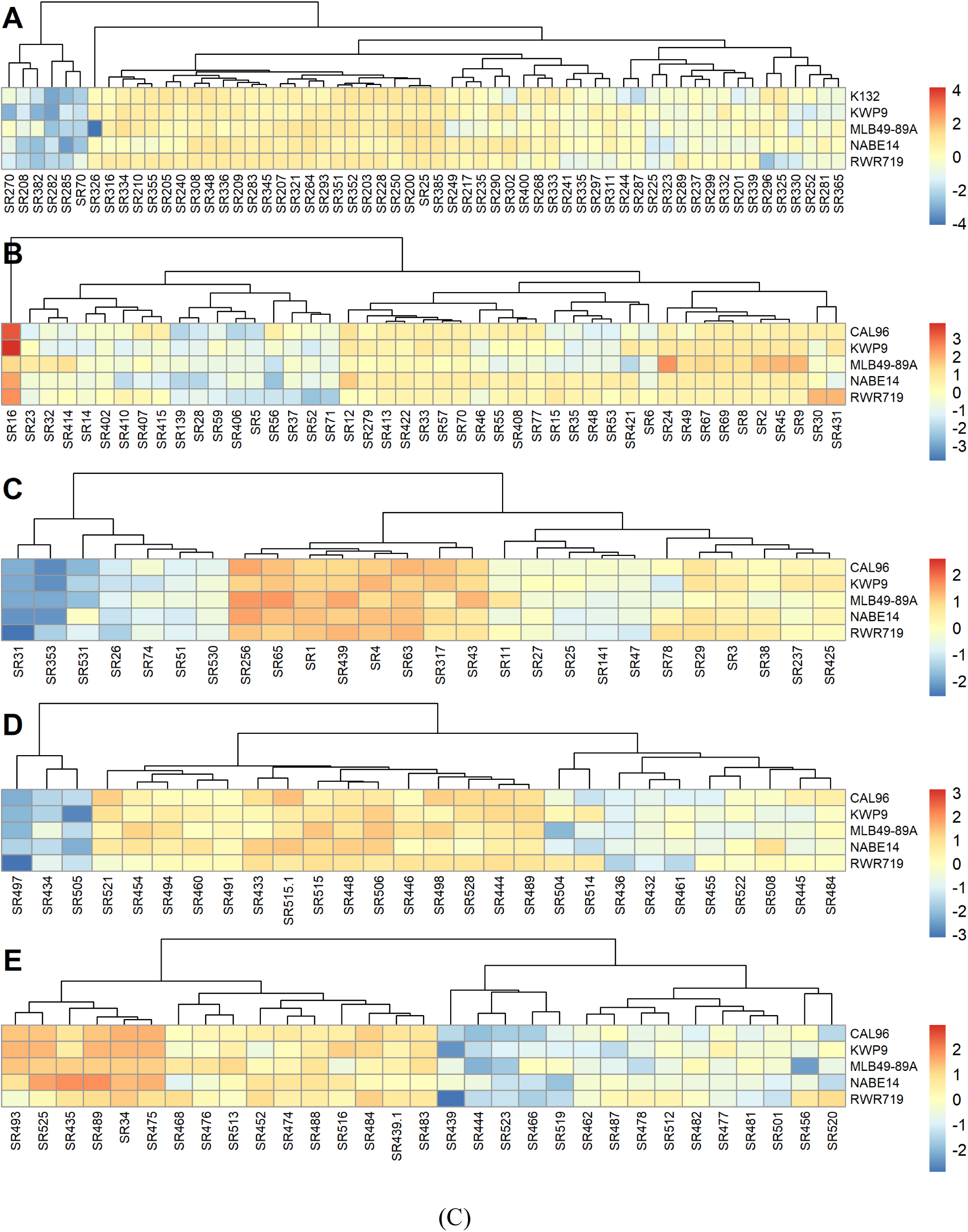
A- Disease severity index for different *S. rolfsii* strains on five common bean varieties across five experiments. B- The heat map of DSI of different *S. rolfsii* isolates. A to E are for experiment 1. 2, 3, 4 and 5 respectively. C- The pre-emergency damping off heat map of different *S. rolfsii* isolates against the five common bean varieties. A to E are for experiment 1, 2, 3, 4 and 5 respectively

**Table 1:** Average DSI of 10 most virulent *Sclerotium rolfsii* strains screened against five common bean varieties in the screen house.

| S/no | Strain | Agro-ecology<sup>1</sup> | District | Year | DSI | SE |
| --- | --- | --- | --- | --- | --- | --- |
| 1 | SR282 | WNMFS | Arua | 2013 | 93.4 | 8.6 |
| 2 | SR417 | TFZ | Bukedea | 2013 | 91.7 | 4.2 |
| 3 | SR406 | LVC | Luweero | 2013 | 84.5 | 2.2 |
| 4 | SR70 | LVC | Mubende | 2013 | 84.3 | 7.8 |
| 5 | SR497 | NMFS | Oyam | 2020 | 82.2 | 1.5 |
| 6 | SR52 | LVC | Luweero | 2013 | 81.8 | 1.5 |
| 7 | SR505 | LVC | Kamuli | 2020 | 80.9 | 0.7 |
| 8 | SR59 | LVC | Mbale | 2013 | 80.6 | 2.9 |
| 9 | SR56 | LVC | Mbale | 2013 | 80.3 | 1.5 |
| 10 | SR410 | LVC | Mbale | 2013 | 80.1 | 1.9 |

<sup>1</sup>WNMFS = West Nile Mixed Farming System, TFZ = Teso farming Zone, LVC = Lake Victoria crescent and Mbale farmland and NMFS = Northern Mixed Farming System.

**Table 2:** Average DSI of 10 least virulent strains when screened against five common bean varieties in the screen house.

| S/no | Strain | Agro-ecology<sup>1</sup> | District | Year | DSI | SE |
| --- | --- | --- | --- | --- | --- | --- |
| 179 | SR45 | LVC | Luweero | 2013 | 18 | 2.9 |
| 180 | SR493 | LVC | Sironko | 2021 | 16.8 | 2.1 |
| 181 | SR435 | WMFS | Hoima | 2021 | 16.1 | 0 |
| 182 | SR454 | WMFS | Hoima | 2021 | 16 | 1.9 |
| 183 | SR264 | LVC | Bugiri | 2013 | 15.9 | 1.5 |
| 184 | SR518 | LVC | Sironko | 2021 | 15.5 | 1.9 |
| 185 | SR506 | LVC | Lwengo | 2020 | 13.2 | 1.5 |
| 186 | SR9 | NMFS | Apac | 2013 | 10.2 | 0.1 |
| 187 | SR489 | LVC | Sironko | 2021 | 10.1 | 1.4 |
| 188 | SR475 | LVC | Sironko | 2021 | 10.1 | 1.1 |

<sup>1</sup>LVC = Lake Victoria crescent and Mbale farm land, WMFS = Western Mixed Farming System and NMFS = Northern Mixed Farming System

When the strains were grouped according to their origin, there was no significant difference in their virulence in experiments 1 (F = 1.25, P = 0.28) and 5 (F = 1.8, P = 0.13). Strains evaluated in experiments 1 and 5 originated from the Eastern Highland, Lake Victoria Crescent and Mbale Farmlands, Northern Mixed Farming System, South Western Highlands, Western Mixed Farming System and West Nile Mixed Farming System agro-ecological zones. However, significant differences were observed in DSI for strains originating from different agro-ecological zones in experiments 2 (F = 8.5, P < 0.0001), 3 (F = 12.2, P < 0.0001) and 4 (F = 5.2, P < 0.0001) (Figure 6). In experiment 2, the DSI of strains from Northern Mixed Farming System was significantly lower than for the rest of the agro-ecological zones. In experiment 3, significant differences in DSI were observed in strains from Teso farming zone, Northern Mixed Farming System and Tanzania while there were no differences in DSI in strains from Lake Victoria Crescent and Mbale Farmland, South Western Highlands and Western Mixed Farming System. In experiment 4, strains from Lake Victoria Crescent and Mbale Farmland and Northern Mixed Farming System varied in DSI while those from Tanzania and Western Mixed farming system did not vary in DSI. Overall, in Uganda, the strains from South Western Highlands had the highest average DSI while strains from Teso Farming Zone had the lowest average DSI. However, strains from Tanzania had the highest average DSI than all the agro-ecological zones of Uganda. When data from all the experiments was pooled, strains from Tanzania and SWH had similar DSI. Similarly, the DSI of strains from EH, NMFS, TFZ and WMFS did not also vary. The strains from LVC and WNMFS also had similar DSI.

**Figure 6:**
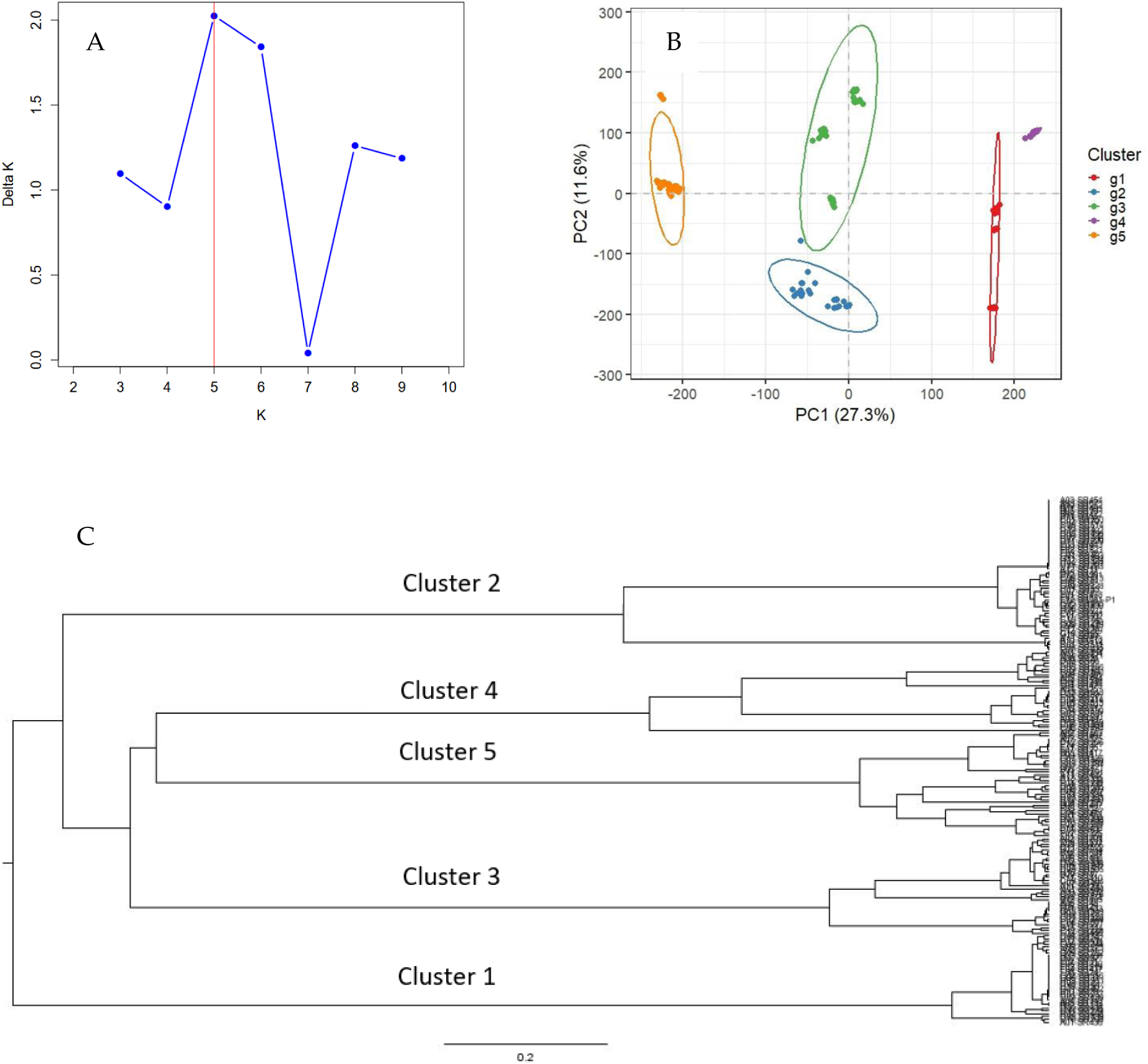
A.- A plot of delta K showing the optimumal number of clusters with a peak at K5.. B- Clustering of *S. rolfsii* isolates using PCA. C- A phylogenetic tree generated using unweighted pair group method with arithmetic mean (UPGMA) based on the two principal components.

**Table 3:**
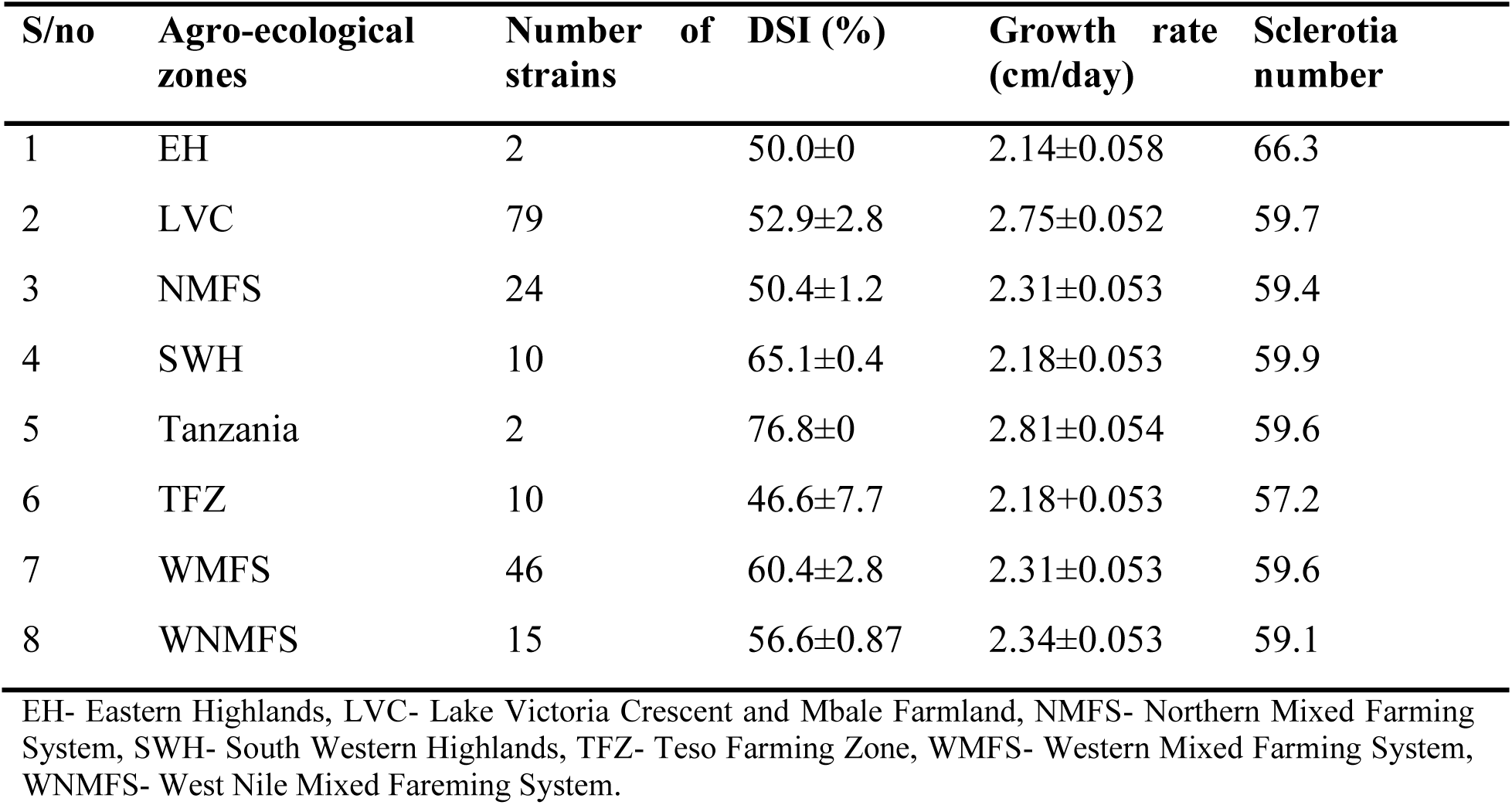
Average disease severity index, growth rate and the number of sclerotia produced by *S. rolfsii* strains from distinct agroecologal zones.

| S/no | Agro-ecological zones | Number of strains | DSI (%) | Growth rate (cm/day) | Sclerotia number |
| --- | --- | --- | --- | --- | --- |
| 1 | EH | 2 | 50.0±0 | 2.14±0.058 | 66.3 |
| 2 | LVC | 79 | 52.9±2.8 | 2.75±0.052 | 59.7 |
| 3 | NMFS | 24 | 50.4±1.2 | 2.31±0.053 | 59.4 |
| 4 | SWH | 10 | 65.1±0.4 | 2.18±0.053 | 59.9 |
| 5 | Tanzania | 2 | 76.8±0 | 2.81±0.054 | 59.6 |
| 6 | TFZ | 10 | 46.6±7.7 | 2.18±0.053 | 57.2 |
| 7 | WMFS | 46 | 60.4±2.8 | 2.31±0.053 | 59.6 |
| 8 | WNMFS | 15 | 56.6±0.87 | 2.34±0.053 | 59.1 |

EH- Eastern Highlands, LVC- Lake Victoria Crescent and Mbale Farmland, NMFS- Northern Mixed Farming System, SWH- South Western Highlands, TFZ- Teso Farming Zone, WMFS- Western Mixed Farming System, WNMFS- West Nile Mixed Farming System.

There was a weak negative correlation between growth rate and pathogenicity (r = -0.06) and number of sclerotia produced (r = -0.02). A weak positive correlation was observed between pathogenicity and number of sclerotia produced (r = 0.06)

#### Susceptibility of common bean varieties to Southern blight

In experiment 1 (F = 2.2, P = 0.07), 2 (F = 1.53, P = 0.19) and 3 (F = 0.21, P = 1.47), there was no significant difference in DSI among the five common bean varieties screened. However, DSI was significantly different among common bean varieties in experiments 4 (F = 7.6 P < 0.0001) and experiment 5 (F= 4.1 p = 0.002) (Figure 5A). A heat map was then generated for DSI caused by the different isolates (Figure 5B). The DSI of RWR719 was the lowest and significantly different from the rest of the varieties in both experiment 4 and 5 while that of NABE14 and KWP9 were different in experiment 4. The DSI of KWP9 and MLB48-89A were different in experiment 5. The DSI of CAL96 and NABE14 were the highest in Experiment 5 and were significantly different from the rest of the varieties. Overall, CAL96 was the most susceptible variety with an average DSI of 52.3% and RWR719 the most resistant with a DSI of 40.5%.

We also assessed seed germination to determine the extent of pre-emergence damping off caused by the different *S. rolfsii* strains (Figure 5C). In all the experiments, there was a significant difference in the effect of the strains on the germination of the five common bean varieties (F = 6.3 P <0.001, F = 8.77 P < 0.001, F = 4.13 P < 0.01, F = 6.87 P < 0.001 and F = 4.55 P < 0.01 for experiments 1, 2, 3, 4 and 5 respectively). Overall, Strain SR31 in experiment 2 from Kisoro district in SWH agro-ecology caused the least germination percentage of 22% while strain SR16 in experiment 2 from Gulu district in NMFS caused the highest germination percentage of 100%. We observed a negative correlation between DSI and germination percentage associated with the isolates (r = -0.58). Isolates that had higher DSI caused lower germination percentage in the common bean varieties

#### 3.4.3 Genetic diversity of *S. rolfsii*

Following sequencing, the libraries generated total reads ranging from 39,134 – 33,014,806 with an average of 6,676,160 reads. The generated reads were mapped to the haploid reference genome assembly for isolate GP3 (Yan et al. 2021. The proportion of mapped reads ranged from 50 to 70% with an average of 60%. Average depth of read per chromosome was 16.5x with a maximum of 83.3 and minimum of 0.06. Out of 188 sequenced isolates, 12 were excluded due to the low depth (< 1x) of sequence coverage. We filtered SNPs for maximum missingness of 15% and minor allele frequency ≥ 0.05 and retained 2,795,782 SNPs. These were pruned based on LD within 50 kb window with 5 kb steps and LD threshold *r*^2^ ≥ 0.5. A set of 51,447 SNPs were retained after LD pruning and used in downstream analyses. . Average minor allele frequency ranged from 0.169 for chromosome 6 to 0.21 for chromosome 1 with an average of 0.209. Average expected heterozygosity (He was 0.782 for all the eight chromosomes while average polymorphic information content (PIC) was 0.322 (Table 4)

**Table 4:**
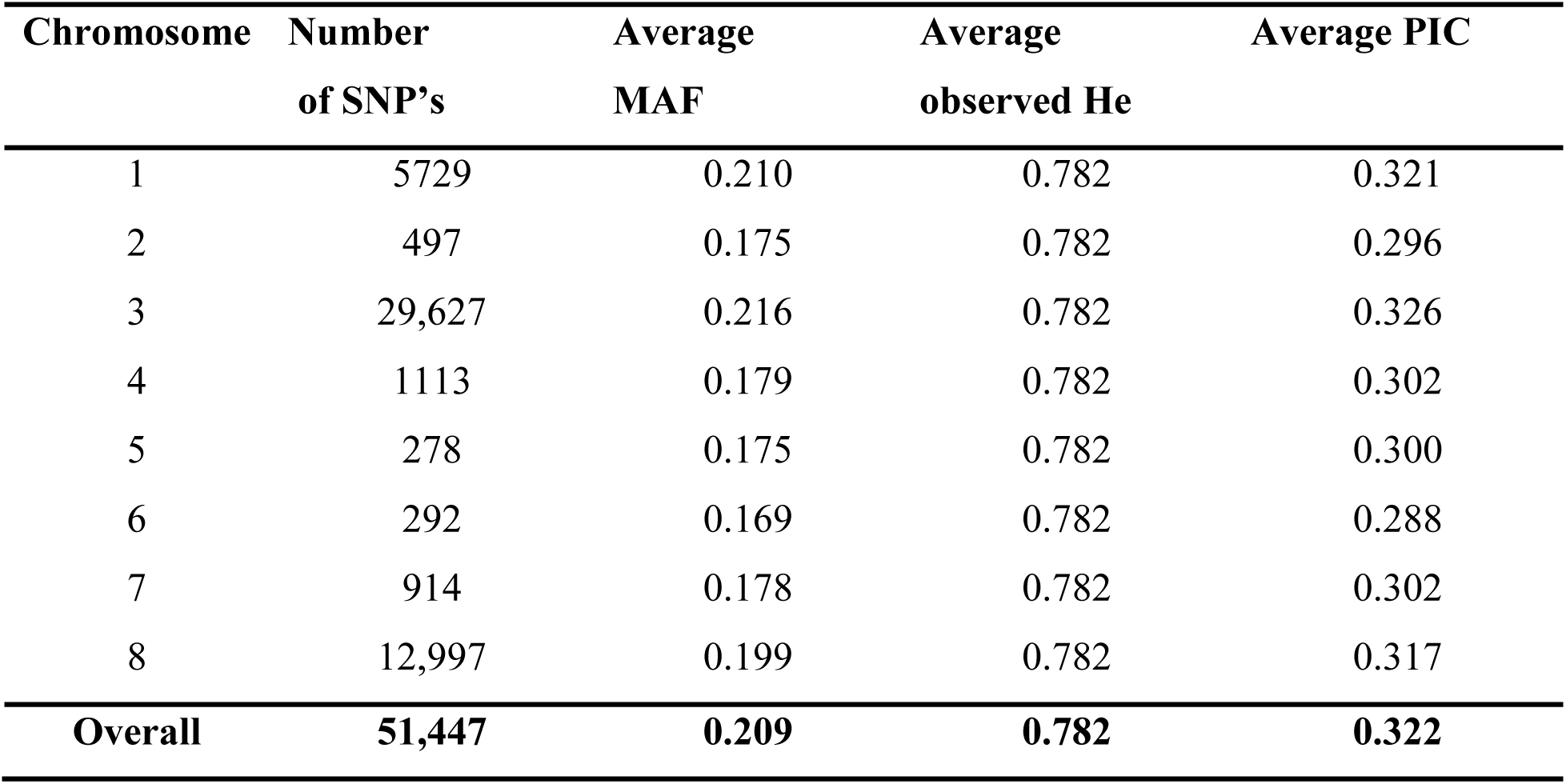
Number of SNPs, avearge minor allele frequency, average expected heterozygosity and average polymorphic information content across all the eight chromosomes.

#### 3.4.3 Population structure and variation of morphological traits between groups

The optimum number of clusters for grouping *S. rolfsii* isolates was determined to be five (Figure 6A and B). Consistent with the results of population structure analyses, a phylogenetic tree had isolates grouped into 5 main branches (Figure 6D).

Admixture analysis was conducted to assess the proportions of ancestry shared among the *S. rolfsii* strains. A total of 44 strains did not have a shared ancestry while 144 strains had a shared ancestry (Table S4). All the strains in cluster 1, 4 and 5 have a shared ancestry while 18 and 26 strains had their ancestry traced entirely traced to cluster 2 and 3 respectively (Table 5) and (Figure 8). A high number of strains from the genetic clusters had their ancestry traced to entirely different genetic clusters. Genetic cluster 1, 2, 3, 4 and 5 had 97, 98, 77, 64 and 55 strains with their ancestry entirely from different genetic clusters. (Table 5 and table S4). When the isolates were grouped into different agro-ecological zones, the 2 isolates frome Tanzania had their ancestry in genetic cluster 2 and 4. Out of the 7 isolates from Teso Farming Zone, 1 isolate had its ancestry entirely in genetic cluster 3 while the rest had a shared ancestry. Of the 43 isolates from Western Mixed Farming System, 6 and 7 isolates had their ancestry entirely in genetic clusters 2 and 3 while the rest has a shared ancestry in the other different genetic clusters. Out of the 14 isolates from West Nile Mixed farming Sytem, 1 isolate had its ancestry entirely in genetic cluster 3 while the rest had a shared ancestry. Of the 26 isolates from Northern Mixed Farming system 2 and 5 isolates had their ancestry entirely in genetic cluster 2 and 3 respectively. For South Western Highlands, 6 isolates had their ancestry entirely in genetic cluster 2. It has been observed that all the isolates that did not have a shared ancestry originated from genetic clusters 2 and 3 (Table S4).

**Table 5:** The number of isolates per genetic cluster and the proportion of ancestry shared among the genetic clusters.

| Genetic cluster | Number of strains with $\phi(\Phi) = 1$ | Number of strains with $\phi(\Phi) = 0$ | Number of strains with $\phi(\Phi) = 0.1-0.9$ |
| --- | --- | --- | --- |
| 1 | 0 | 97 | 91 |
| 2 | 18 | 98 | 72 |
| 3 | 26 | 77 | 85 |
| 4 | 0 | 64 | 124 |
| 5 | 0 | 55 | 133 |

When the K values were varied from 2 to 5 according to procedure by Kopelman et al. (2015), all the isolates from Lake victoria Crescent and Mbale farmland and Western Mixed Farming Sytem were present in all the genetic clusters for the different K values. Meanwhile, some isolates from South Western Highland, Northern Mixed Farming System, Eastern Highlands, West Nile Mixed Farming System, Teso Farming Zone and Tanzania were missing in some clusters for the different K values. (Table 6). There were only two isolates from EH and Tanzania and they clustered in a single genetic cluster though the K values were beign increased from K2 to K5.

**Table 6:**
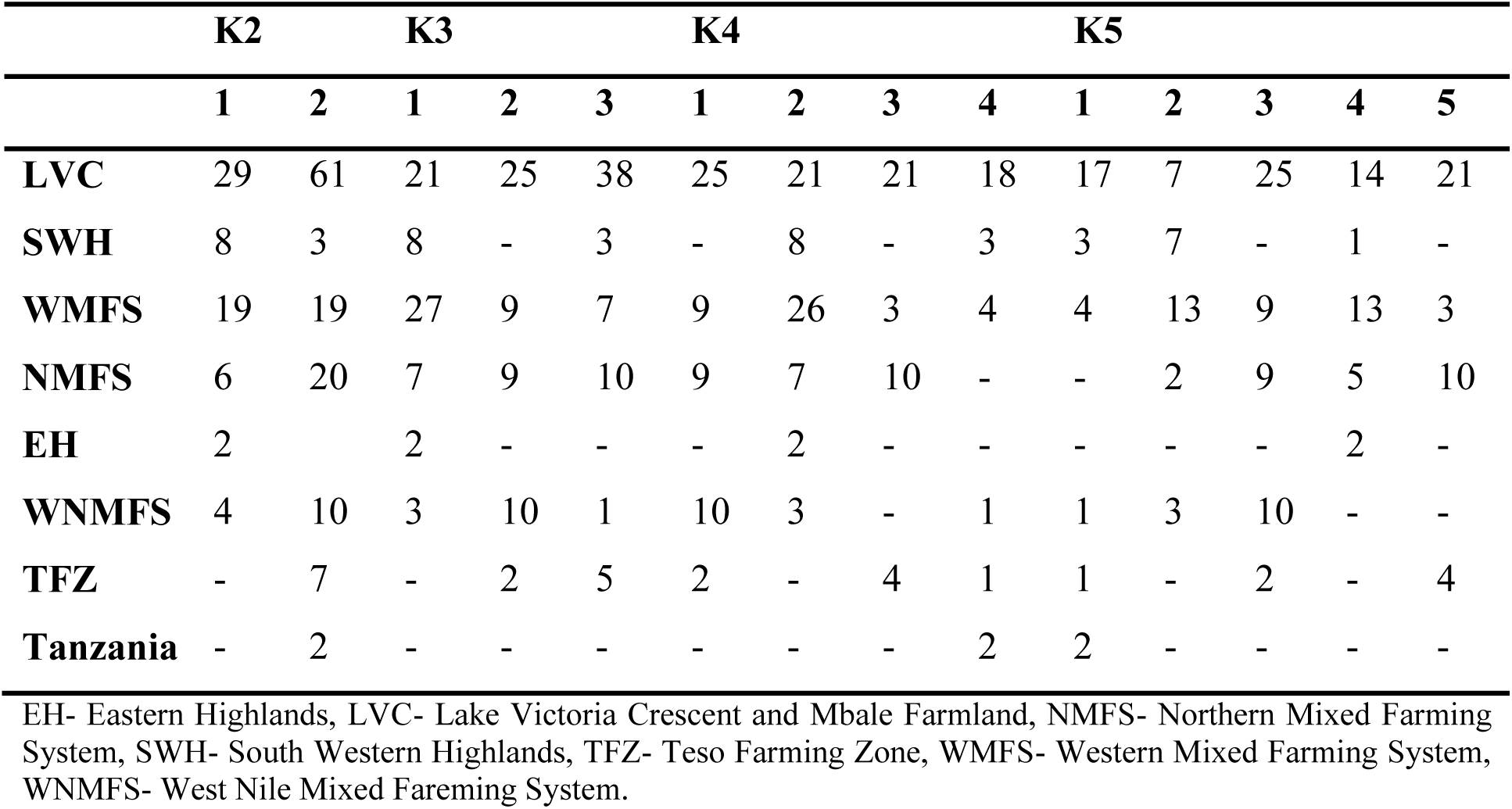
The number of isolates from the different agro-ecological zones in the different genetic clusters when the K values were varied from K2 to K5.

Analysis of variance was conducted to compare variation in the morphological and virulence traits among these five genetic clusters. There were significant differences in DSI caused by strains from different genetic clusters (F = 9.102, P = 0.001) (Figure 7A). The DSI of strains from cluster 4 was significantly different from that of the other clusters. The growth rate of strains in different clusters also varied significantly (F = 6.5, P = 0.001). The strains from clusters 1, 2 and 3 had similar growth rate while the growth rate of strains from cluster 4 and 5 differed significantly (Figure 7B and table S5). The number of sclerotia produced by the strains from different clusters were significantly different too (F = 8.9, P = 0.001). The number of sclerotia produced by strains from clusters 1, 4 and 5 were significantly different while the number of sclerotia produced by strains in cluster 2 and 3 did not vary (Figure 7C and table S3). There was a significant association between the genetic clusters and number of sclerotia (X^2^ = 834.3, P= 0.00001), growth rate (X^2^ = 639, P= 0.000013) and DSI (X^2^= 1346, P= 0.00001).

Nei genetic distance was used to assess the genetic differentiation among isolates from different agro-ecological zones. The genetic distance ranged from 0.001 between Lake Victoria Crescent and Mbale farmland and Northern Mixed Farming System to 0.089 between Teso Farming Zone and West Nile Mixed Farming System (Table 7A). The genetic identity of the isolates was assessed among the different agro-ecological zones. The similarity coefficient ranged from 0.887 between isolates from Bukoba in Tanzania and West Nile Mixed Farming System to 0.999 between Eastern Highland and South Western Highland. (Table 7B). The average genetic distance across all the agro-ecological zone was 0.038 while the average genetic identity across all the agro-ecological zones was 0.963.

**Figure 7:**
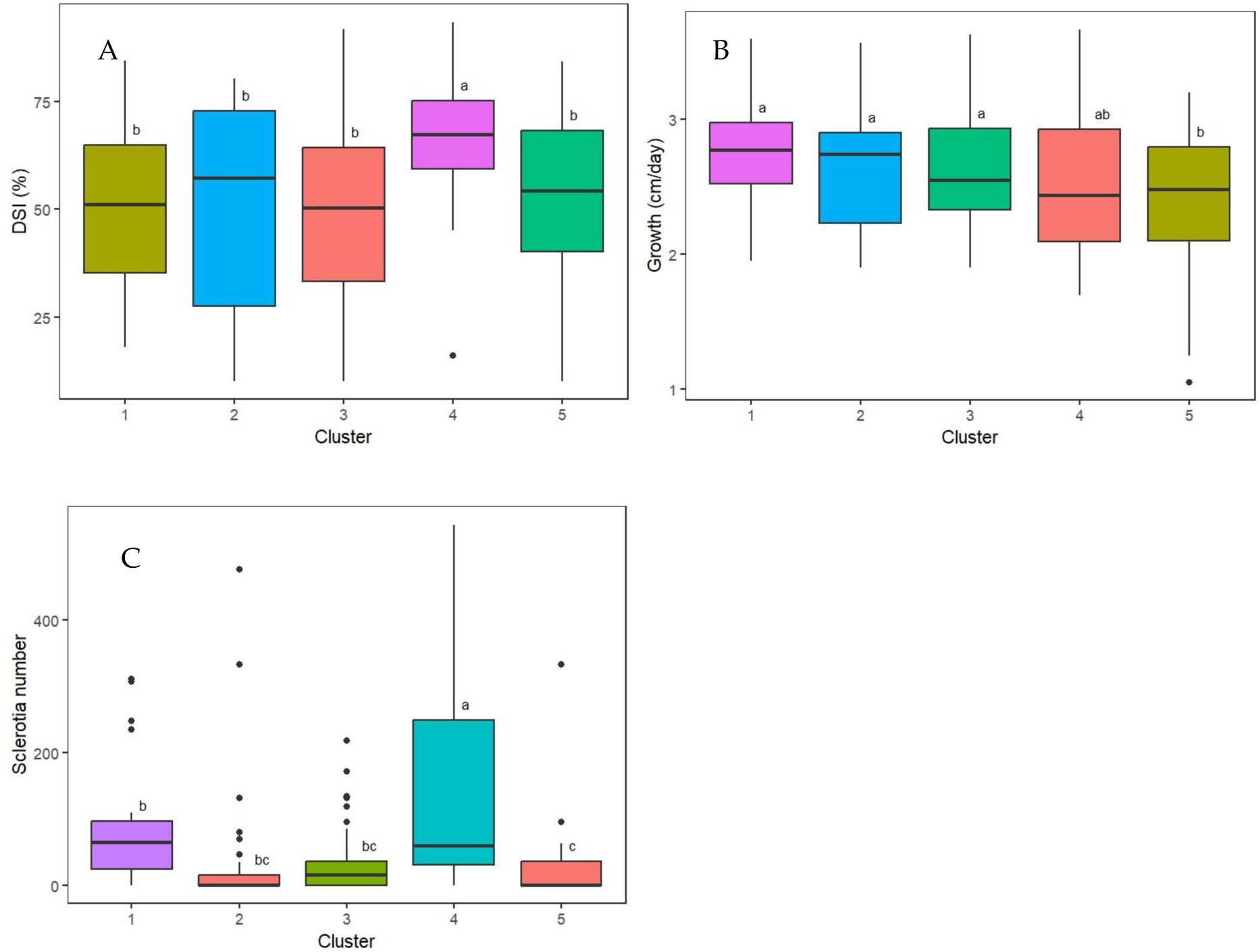
Comparison of morphological and pathogenicity characteristics of *S. rolfsii* isolates among five genetically distinct clusters. A. Variation in DSI. B. Number of sclerotia produced by strain from different groups. C. Growth rate of the strains in the five genetic groups

**Figure 8.**
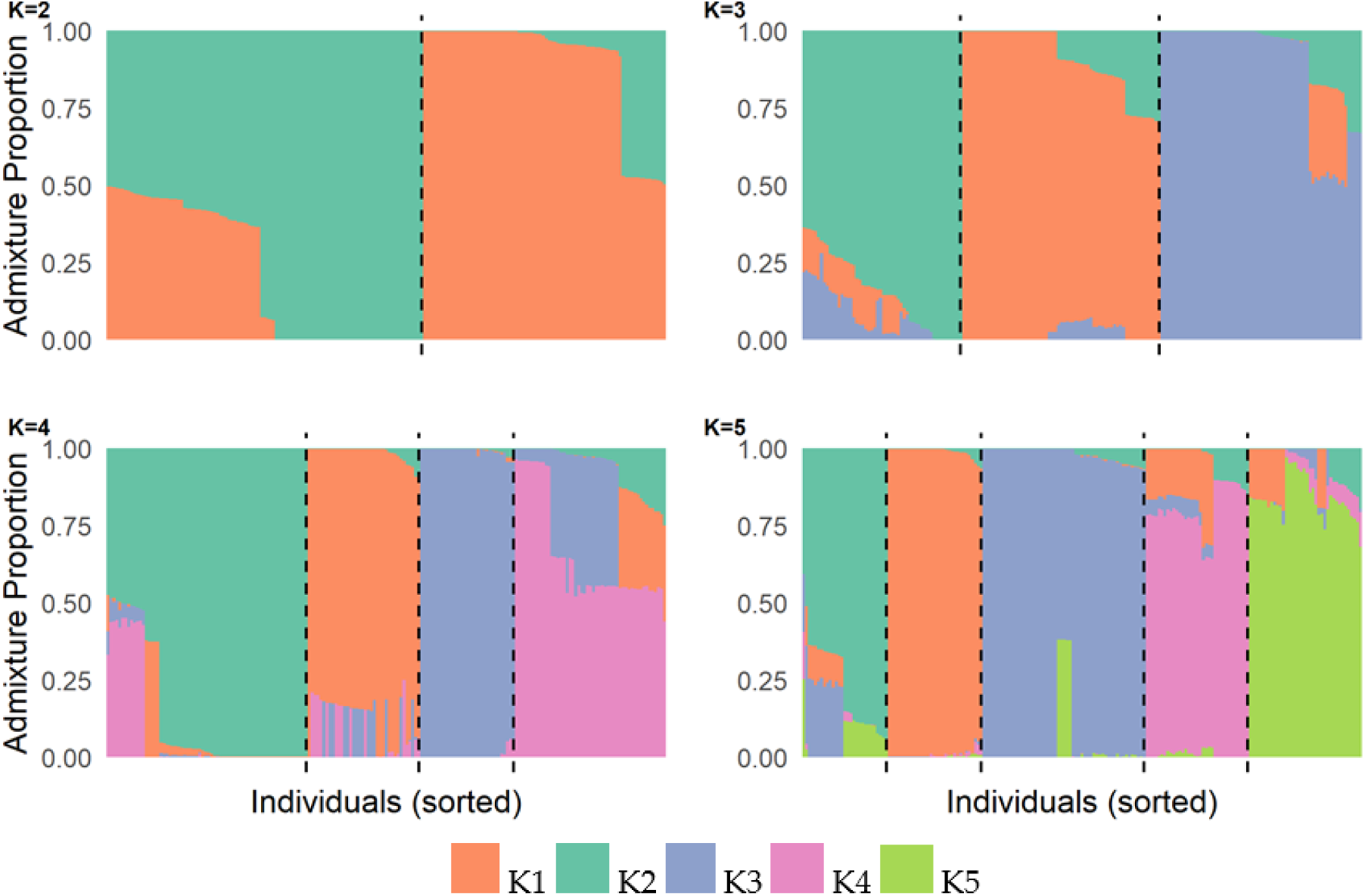
– Proportions of *S. rolfsii* isolates ancestry assigned to K populations. (I suggest to make figure with K ranging from 2 to 5 and group isolates by their geographic origin. This arrangement of data will give enough information to start discussing population structure. The plot below is not very helpful…. Optimal number of cluster has no much meaning usually, unless fit to some biological or geographical patterns. Best is to start with K = 2 and check does the grouping of isolates at K = 2 explains anything about geography of biology of isolates? Then move to K = 3 and so on until to K = 5. Main goal is to try to explain biology and geography by clustering at different values of K. Use newly arranged plot and Q matrix data generated by Admixture to discuss the proportions of ancestry shared between groups of isolates.

**Table 7: A.**
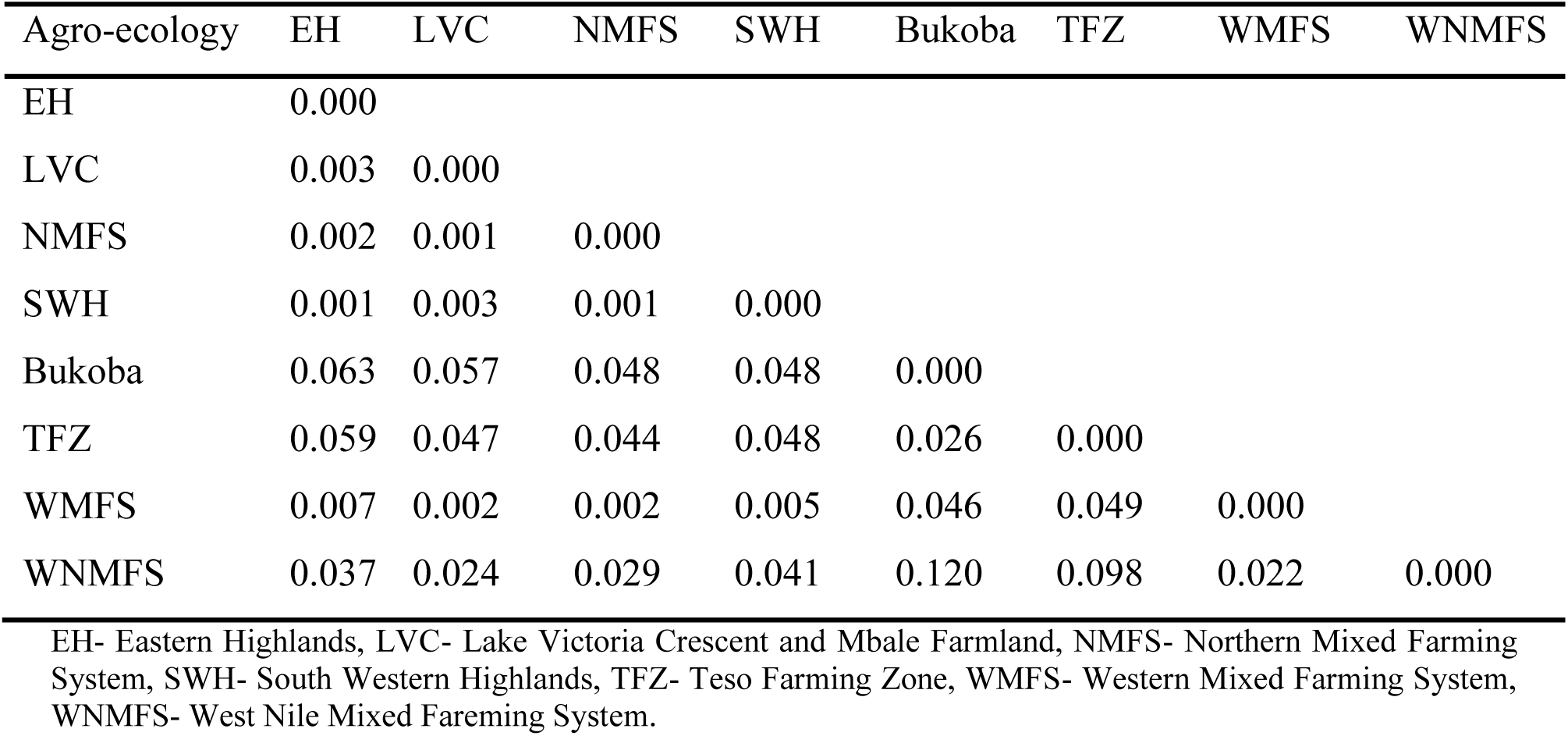
– Pairwise comparison of genetic distance between isolates from different agro-ecological zones.

**Table 7: B.** – The similarity coefficient of isolates from different agro-ecological zones.

| Agro-ecology | EH | LVC | NMFS | SWH | Bukoba | TFZ | WMFS | WNMFS |
| --- | --- | --- | --- | --- | --- | --- | --- | --- |
| EH | 1.000 |  |  |  |  |  |  |  |
| LVC | 0.997 | 1.000 |  |  |  |  |  |  |
| NMFS | 0.998 | 0.999 | 1.000 |  |  |  |  |  |
| SWH | 0.999 | 0.998 | 0.999 | 1.000 |  |  |  |  |
| Bukoba | 0.939 | 0.945 | 0.953 | 0.953 | 1.000 |  |  |  |
| TFZ | 0.943 | 0.954 | 0.957 | 0.953 | 0.975 | 1.000 |  |  |
| WMFS | 0.993 | 0.998 | 0.998 | 0.995 | 0.955 | 0.952 | 1.000 |  |
| WNMFS | 0.963 | 0.976 | 0.971 | 0.960 | 0.887 | 0.907 | 0.978 | 1.000 |

EH- Eastern Highlands, LVC- Lake Victoria Crescent and Mbale Farmland, NMFS- Northern Mixed Farming System, SWH- South Western Highlands, TFZ- Teso Farming Zone, WMFS- Western Mixed Farming System, WNMFS- West Nile Mixed Farming System.

Analysis of molecular variance (AMOVA) was conducted to assess proportions of genetic variance within and among groups of isolates defined by their genetic relatedness or origin in distinct agro-ecological zones (Table 8). Results revealed that 0.39% of the variation was between population while 11.98% was between samples within population and 87.62% between samples across populations. The low proportion of variance explained by groups indicates the lack of strong population structure in the sampled collection of *S. rolfsii* isolates.

**Table 8:** AMOVA outputs based on 5 optimal genetic clusters and for the 8 different geographical locations.

| **AMOVA based on the five optimal genetic clusters** |  |  |  |  |  |
| --- | --- | --- | --- | --- | --- |
| **Sources of variation** | **Df** | **SS** | **MSS** | **%** | **Phi** |
| Between populations (Pop) | 4 | 82337.64 | 20584.41 | 0.39 | 0.123 |
| Between samples within pop | 184 | 3035249.01 | 16495.92 | 11.98 | 0.120 |
| Within samples | 189 | 2448136.00 | 12953.10 | 87.62 | 0.003 |
| Total | 377 | 5565722.65 | 14763.19 | 100 |  |
| **AMOVA among agro-ecological zones** |  |  |  |  |  |
| **Sources of variation** | **Df** | **SS** | **MSS** | **%** | **Phi** |
| Between agro-ecology | 8 | 35,999 | 4,499 | 0 | 0.122 |
| Within agro-ecology | 180 | 1,209,370 | 6,718 | 100 | 0.001 |
| Total | 188 | 1,245,369 | 6,624 | 100 |  |

The 11.98% of variance explained by between samples within populations suggest that distinct populations experience different levels of intermating among *S. rolfsii* isolates, though these differences are not substantial. The high proportion of genetic variance attributed to within sample level suggests extensive level of intermating among *S. rolfsii* isolates in the collection. Analysis of molecular variance conducted on isolates grouped based on the agro-ecological zone of origin reveals no genetic differentiation among samples collected at different locations.

## Discussion

### Morphological characterization of *S. rolfsii* strains

We have characterized 84 newly collected strains of *S. rolfsii* not previously characterized by Paparu et al. (2020) for growth rate and number of sclerotia produced when grown on Potato Dextrose Agar. *S. rolfsii* is known for its prolific growth rate *in vitro*. In the current study, growth rate ranged from 1.1 to 3.6 cm per day on PDA at 25 °C. These estimates are within the ranges reported in the previous studies. Paparu *et al*. (2020) obtained growth rate of 1.04 cm to 2.92 cm per day on PDA, while growth rates reported by Le *et al*. (2012) ranged from 0.67 to 1.8 cm per day. Paul et al. (2023) studied the growth rate of *Athelia rolfsii* causing stem rot in peanuts. A high growth rate of 3.6 cm per day was reported on PDA while lower growth rates were reported on oatmeal agar and corn meal agar. The growth rate of 66 *Athelia rolfsii* Isolates in China ranged from 1.06 to 2.06 cm/day (Yan et al. 2022). Paul *et al*. (2023) showed that growth rates depend on temperature, with the highest and lowest growth rates observed at 30 **°C** and 15 **°C**, respectively.

In the current study, there was a weak negative correlation between growth rate and pathogenicity and number of sclerotia produced. While Yan *et al*. (2022) reported lack of significant differences in the growth rate of a highly and weakly aggressive strains of *S. rolfsii* causing peanut stem rot, other studies showed a positive correlation between pathogenicity and number of sclerotia produced (Yan et al. 2022; Zamini-Noor et al. 2024). Similarly, Paparu *et al*. (2020) also reported moderate correlations in their study.

Comparison of the number of sclerotia produced in different studies show that *S. rolfsii* strains are capable of producing a high number of sclerotia. In peanuts, 35 strains produced less than 100 sclerotia per petri dish while 25 strains produced more than 300 (Yan *et al*., 2022). In our study, 75 strains did not produce sclerotia by day 28 in the study of Xie *et al*. (2014) only one strain out of 19 studied did not produce sclerotia. However, we observed that when the cultures were kept for more than two months, all the strains eventually produced sclerotia, suggesting that all strains of *S. rolfsii* are likely able to produce sclerotia under conditions of nutrient stress. This is compatible with the fact that the sclerotia is known to be a fruiting body that is used by the pathogen for transition from one season to another, even in the absence of the host plant (Agrios 2005).

### Pathogenicity of *S. rolfsii* strains on Common bean

We carried out pathogenicity test on 188 *S. rolfsii* strains using five common bean varieties. Importantly, we found that strains stored for over 10 years were still virulent on common bean. The ability to store strains for long periods of time opens opportunities for creating collections of strains, a resource that is critical for 1) improving pathogen surveillance system, 2) studying pathogen population dynamic over time, 3) studying pathogen adaptation to environmental changes, and 4) conducting virulence studies to identify resistance crop varieties. All analyzed *S. rolfsii* strains were virulent on common bean and caused all the symptoms associated with southern blight such as brown water-soaked lesions at the collar of seedling, presence of white cottony mycelia on lesions, wilting of plants, presence of sclerotia at the collar of plants and pre-emergence and post-emergence dumping off. Some strains caused severe disease while others caused mild disease. Similarly, the strains also differed in their effect on germination of common bean varieties, with some strains causing more pre-emergence damping off than others. The high Disease Severity Index (DSI) was associated with the high frequency of pre-emergence damping off. The agro-ecological zones with most pathogenic strains were Northern Mixed Farming System, South Western Highlands, West Nile Mixed Farming System, Lake Victoria Crescent and Mbale Farmlands, Western Mixed Farming System, Eastern Highlands and Teso Farming Zone respectively. The strains from Bukoba in Tanzania had highest average DSI compared to the ones from Ugandan agro-ecological zones. This observation may have serious implications for the spread of these highly virulent strains to other regions of East Africa, because traders often transport grains across locations and farmers use these grains as seed sources. Moreover, since *Fusarium* species can also undergo sexual reproduction with compatible thalli, movement of these virulent strains may increase virulence of Ugandan strains if their strains mate.

In the current study, DSI of the analyzed strains ranged from 10.1 to 93.3%, which is comparable to the previous report by Paparu *et al*. (2020), where DSI ranged from 4.4 to 100%. The DSI caused by different strains on common bean also varied across varieties, with RWR719 being the most resistant and CAL96 being the most susceptible variety. Similarly, in the study by Paparu et al. (2020), RWR719 was also the most resistant variety while CAL96 was the most susceptible. Other studies have reported variation in the pathogenicity of *S. rolfsii* strains in various crop varieties (Narayan et al. 2018; Modal et al. 2022: Parvin et al. 2016). Narayan *et al*. (2018) reported variations in the virulence of *S. rolfsii* on different sweet potato varieties, concluding that different races of *S. rolfsii* might exist in South Korean.

*S. rolfsii* has a broad host range and causes disease in various crops. Paparu *et al*. (2020) obtained a strain HOI356-Peanut from a volunteer groundnut plant in the bean field that was also pathogenic on common bean. In the current study, the name was changed to SR411 and it caused a DSI of 34.5%. In the pathogenicity studies of *S. rolfsii* obtained from groundnuts, tomatoes and taro, strains from tomatoes and taro were found to be pathogenic on groundnuts (Le *et al*., 2012). Nineteen *S. rolfsii* strains collected from various crops in Southern United States (Florida, Georgia, Louisiana, South Carolina, Texas, and Virginia) were also pathogenic on tomatoes, peppers and peanuts (Xie *et al*., 2014). Paul *et al*. (2023) used two *S. rolfsii* strains from common bean to conduct pathogenicity on other crops and showed that both strains were pathogenic on tomatoes, eggplant and chickpea but not on chili, soybean and cowpea. In spite of a wide host range, not all strains are pathogenic across all *S. rolfsii* hosts. Resource-constrained farmers need to implement appropriate crop rotation strategies by including non-host crops if they are to manage the southern blight through crop rotation to reduce inoculum levels in the soil.

### Molecular diversity of *S. rolfsii* strains

In this study, the expected heterozygosity and polymorphic information content of *S. rofsii* strains was high, implying a high level of genetic diversity among the collected strains. This result therefore implies that larger collection of diverse strains need to be used for representative screening of germplasm for disease resistance. This can increase the cost of screening process. While studying the genetic diversity of *S. rolfsii* infecting various crops in China using RAPD, Wang et al. (2023) observed a polymorphic information content of greater than 0.5. Meena et al. (2023) observed a polymorphic information content of 0.242 to 0.536 among S. rolfsii infecting groundnuts while using RAPD. However, it is difficult to compare the finding in these studies with the current study because they used different methodologies.

Analysis of population structure indicates that strains could be optimally clustered into 5 genetically distinct groups. The population of *S. rolfsii* from the Ugandan agro-ecological zone showed evidence of admixture with the strains from other agro-ecological zones. The strains from these five genetic groups showed significant differences in DSI, number of sclerotia produced and the rate of growth. It is not immediately clear whether these differences have adaptive value or not, because no strong correlation between virulence of common bean and these morphological and pathogenicity characteristics of strains was detected. Meanwhile, in the study by Yan *et al*. (2022) while clustering strains of *Athelia rolfsii* causing stem rot in peanuts in China using ISSR markers, the strain clustered based on geographical origin and sclerotia traits but not virulence. In the current study, the strains did not cluster into the genetic groups by agro-ecological zone of origin. Contrary to these results, clustering of *S. rolfsii* strains pathogenic to groundnuts using sequence data from the ITS of ribosomal DNA was consistent with the strains’ geographic origins (Le et al., 2012).

The genetic distance between the isolates from the different agro-ecological zones was 0.038 while the average similarity coefficient between the isolates from different agro-ecological zones was 0.968. These results indicate that the isolates are closely related and have a recent common ancestry. Meena et al. (2023) used ITS and RAPD primers to study genetic diversity among *S. rolfsii* infecting groundnut. In the study, similarity coefficient ranged from 0.33 to 0.57 between different isolates. While using RAPD markers to studt genetic diversity among *S. rolfsii* causing collar rot in Chick pea, Srividya et al. (2022) observed a similarity index of 0.15 to 0.72 and disimilarity index of 0.75 to 0.83. The concluded that the populations of this fungus are made up of more than one strain and that few are derived clonally.

Following AMOVA, 11.98% of the genetic variation was within strains within population, while 87.62% was within strains across population and only 0.39% was between populations. When strains were grouped based on the agro-ecological zones of origin, 100% of the variation was within agro-ecological zones and no variation explained by geographic origin. These results indicate very low levels of genetic differentiation among populations, which could be associated with substantial inter-population gene flow or shared ancestry. The moderate values of genetic variance between populations (φ = 0.1238) and between samples within populations (φ = 0.1203 and φ = 0.1231) suggest that most genetic variance is within populations rather than between populations. This result could indicate that there are high levels of inter-mating among isolates within the sampled geographic range or there is long-distance movement of isolates. The latter could be facilitated by the widespread pulse movement across the country, as farmers often purchase them and use as seed for the next planting season. While analysing population structure of *S. rolfsii* strains infecting Chinese herbal crops, Wang *et al*. (2023) identified three groups and the differences between the groups was mostly related to geographical location. The analysis of molecular variance showed that 28% of the variation was among the groups while 72% was within groups. In their study, Cluster analysis showed that the closely spaced populations grouped in one cluster, this implies that the isolates in their study may not have been highly mobile. In the study by Mehri et al. (2020), 70% and 30% of the observed variance corresponded to the difference between and within the populations, respectively. They suggested that mating between populations would be less likely and thus, gene flow is restricted.

## Conclusion and recommendations

*Sclerotium rolfsii* causing southern blight of common bean is a morphologically and genetically diverse pathogen. Significant differences in morphological and virulence traits have been detected among genetically differentiated groups of isolates. The collected *S. rolfsii* strains showed high levels of genetic diversity. However, quantifying genetic variance at different levels of population hierarchy showed that most of genetic variance is explained by variation within isolates and between isolates within populations. Our study shows the low level of population stratification in *S. rolfsii* and lack of strong genetic differentiation among isolates from distinct geographic regions. The high levels of genetic diversity in *S. rolfsii* suggest that multiple isolates might be required for adequate screening of germplasm to identify novel sources of resistance. Since there are no regional specific virulent isolates to common bean, new resistant varieties developed and evaluated in one region could easily adapt to other regions within the country, reducing the burden to breed region-specific varieties.”

The movement of common bean seeds across Uganda and the re-use of these seeds is subsequent production seasons by farmers likely had a significant impact on the genetic diversity of *S. rolfsii* population and likely it was the key factor promoting inter-mating among different populations. To prevent the introduction of virulent strains from Tanzania to Uganda, better control of germplasm movement needs to be implemented.

## Acknowledgements

We are grateful to the staff of Legumes Program at the National Crops Resources Research Institute, Namulonge, Kansas State University, Department of Plant Pathology, Makerere University Regional Center for Crop Improvement (MaRCCI). This work was supported by the Bill and Melinda Gates Foundation (INV-004430).

## Notes

### Competing Interest Statement

The authors have declared no competing interest.

