## Supplementary material for "Pathogenic and genetic diversity of *Sclerotium rolfsii,* the causal agent of Southern blight of common bean in Uganda": Table S1.docx

**Table S1:** List of S. rolfsii isolates, their agroecology of origin, virulence and genetic cluster

| S/no | Strain | Agroecology | District | DSI | SE | Genetic cluster |
| --- | --- | --- | --- | --- | --- | --- |
| 1 | SR282 | WNMFS | Kabale | 93.4 | 8.6 | 4 |
| 2 | SR417 | TFZ | Bukedea | 91.7 | 4.2 | 3 |
| 3 | SR406 | LVC | Luweero | 84.5 | 2.2 | 1 |
| 4 | SR70 | LVC | Mubende | 84.3 | 7.8 | 5 |
| 5 | SR497 | NMFS | Oyam | 82.2 | 1.5 | 3 |
| 6 | SR52 | LVC | Luweero | 81.8 | 1.5 | 3 |
| 7 | SR505 | LVC | Kamuli | 80.9 | 0.7 | 1 |
| 8 | SR59 | LVC | Mbale | 80.6 | 2.9 | 4 |
| 9 | SR56 | LVC | Mbale | 80.3 | 1.5 | 2 |
| 10 | SR410 | LVC | Mbale | 80.1 | 1.9 | 1 |
| 11 | SR523 | WMFS | Hoima | 80 | 0.9 | 5 |
| 12 | SR208 | LVC | Sironko | 79.2 | 7.3 | 4 |
| 13 | SR74 | LVC | Nakaseke | 78.6 | 1.1 | 5 |
| 14 | SR26 | SWH | Kabale | 78.5 | 3.4 | 2 |
| 15 | SR48 | LVC | Luweero | 78.3 | 0.1 | 2 |
| 16 | SR37 | WMFS | Kyenjojo | 78 | 0 | 4 |
| 17 | SR531 | TANZANIA | Bukoba | 77.7 | 0.8 | 2 |
| 18 | SR439 | WMFS | Hoima | 77.3 | 0 | 4 |
| 19 | SR35 | NMFS | Kitgum | 77 | 2.3 | 3 |
| 20 | SR530 | TANZANIA | Bukoba | 76.5 | 0 | 2 |
| 21 | SR402 | SWH | Kisoro | 75.8 | 0.1 | 4 |
| 22 | SR6 | LVC | Apac | 75.2 | 3.6 | 3 |
| 23 | SR51 | LVC | Luweero | 75.2 | 3 | 3 |
| 24 | SR28 | SWH | Kabale | 75.2 | 0.3 | 4 |
| 25 | SR225 | WNFS | Arua | 75 | 6.9 | 4 |
| 26 | SR78 | LVC | Nakaseke | 74.7 | 0 | 5 |
| 27 | SR25 | SWH | Kabale | 74.5 | 1.9 | 2 |
| 28 | SR335 | NMFS | Oyam | 74.3 | 6.9 | 3 |
| 29 | SR323 | Unknown | Unknown | 74.2 | 6.9 | 5 |
| 30 | SR32 | LVC | Kamuli | 73.9 | 2 | 5 |
| 32 | SR510 | WNFS | Arua | 72.3 | 0.9 | 2 |
| 33 | SR237 | LVC | Lwengo | 71.7 | 1.6 | 4 |
| 34 | SR438 | WMFS | Hoima | 71 | 3.6 | 4 |
| 35 | SR415 | LVC | Mukono | 70.9 | 1.1 | 5 |
| 36 | SR466 | LVC | Sironko | 70.8 | 0.9 | 3 |
| 37 | SR471 | LVC | Sironko | 70.8 | 3.7 | 3 |
| 38 | SR53 | LVC | Mbale | 70.7 | 0.1 | 1 |
| 39 | SR38 | WMFS | Kyenjojo | 70.4 | 2 | 5 |
| 40 | SR407 | LVC | Luweero | 70.4 | 1.2 | 3 |
| 41 | SR425 | LVC | Sironko | 70.2 | 0 | 3 |
| 42 | SR65 | LVC | Mukono | 69.8 | 1.4 | 5 |
| 43 | SR46 | LVC | Luweero | 69.5 | 0.9 | 5 |
| 44 | SR24 | SWH | Kabale | 69.5 | 4.8 | 4 |
| 45 | SR421 | LVC | Luweero | 69.1 | 0.2 | 3 |
| 46 | SR77 | LVC | Nakaseke | 68.8 | 0.1 | 2 |
| 47 | SR434 | WMFS | Hoima | 68.6 | 0.6 | 3 |
| 48 | SR11 | NMFS | Apac | 68.2 | 4.7 | 3 |
| 49 | SR30 | SWH | Kisoro | 68 | 1.6 | 4 |
| 50 | SR55 | LVC | Mbale | 67.9 | 2.8 | 5 |
| 51 | SR15 | NMFS | Gulu | 67.7 | 1.1 | 3 |
| 52 | SR436 | WMFS | Hoima | 67.7 | 3.7 | 4 |
| 53 | SR139 | LVC | Sironko | 67.3 | 0 | 4 |
| 54 | SR141 | LVC | Wakiso | 67.2 | 0.6 | 3 |
| 55 | SR519 | WMFS | Hoima | 67.1 | 3.8 | 4 |
| 56 | SR12 | NMFS | Apac | 67 | 2.5 | 1 |
| 57 | SR279 | WNFS | Arua | 66.3 | 1.8 | 5 |
| 58 | SR5 | TFZ | Amuria | 65.2 | 0 | 5 |
| 59 | SR299 | WMFS | Ibanda | 64.9 | 6 | 3 |
| 60 | SR201 | EH | Kapchorwa | 64.8 | 6 | 1 |
| 61 | SR47 | LVC | Luweero | 64.8 | 2.6 | 1 |
| 62 | SR330 | NMFS | Oyam | 64.3 | 6 | 4 |
| 63 | SR241 | LVC | Masaka | 64.3 | 5.9 | 3 |
| 64 | SR509 | WNFS | Arua | 63.5 | 1.6 | 5 |
| 65 | SR281 | WNFS | Arua | 63.4 | 5.9 | 2 |
| 66 | SR23 | SWH | Kabale | 63.3 | 0 | 4 |
| 67 | SR527 | WMFS | Hoima | 63.3 | 3.2 | 5 |
| 68 | SR478 | LVC | Sironko | 63 | 2.3 | 5 |
| 69 | SR296 | LVC | Rakai | 62.2 | 5.8 | 3 |
| 70 | SR482 | LVC | Sironko | 61.9 | 0 | 1 |
| 71 | SR449 | WMFS | Hoima | 61.7 | 2.1 | 2 |
| 72 | SR14 | WNFS | Arua | 61.6 | 0 | 2 |
| 73 | SR27 | SWH | Kabale | 61.6 | 0.3 | 2 |
| 74 | SR400 | LVC | Mbale | 61.4 | 5.7 | 1 |
| 75 | SR356 | NMFS | Lira | 61.3 | 5.7 | 4 |
| 76 | SR422 | LVC | Kayunga | 61.2 | 0.2 | 3 |
| 77 | SR339 | NMFS | Oyam | 61.2 | 5.7 | 3 |
| 78 | SR413 | LVC | Kayunga | 60.8 | 1.1 | 5 |
| 79 | SR520 | WMFS | Hoima | 60.8 | 0 | 3 |
| 80 | SR244 | Unknown | Unknown | 60 | 5.6 | 4 |
| 81 | SR287 | WMFS | Mbarara | 59.7 | 5.5 | 1 |
| 82 | SR464 | LVC | Sironko | 59.7 | 3.9 | 4 |
| 83 | SR514 | WNFS | Arua | 59.5 | 0 | 5 |
| 84 | SR297 | WMFS | Kabarole | 59.3 | 5.5 | 4 |
| 85 | SR31 | SWH | Kisoro | 58.2 | 0 | 4 |
| 86 | SR456 | LVC | Sironko | 58.1 | 0 | 5 |
| 87 | SR2 | TFZ | Amuria | 58 | 0 | 2 |
| 88 | SR501 | NMFS | Oyam | 58 | 3.7 | 1 |
| 89 | SR455 | WMFS | Hoima | 57.8 | 0 | 5 |
| 90 | SR487 | LVC | Sironko | 57.6 | 1.7 | 3 |
| 91 | SR207 | LVC | Jinja | 57.4 | 5.3 | 5 |
| 92 | SR250 | LVC | Rakai | 56.4 | 5.2 | 3 |
| 93 | SR332 | NMFS | Oyam | 56.1 | 5.2 | 5 |
| 94 | SR499 | NMFS | Oyam | 56.1 | 0 | 3 |
| 95 | SR477 | LVC | Sironko | 55.6 | 0 | 1 |
| 96 | SR57 | LVC | Mbale | 55.6 | 4 | 3 |
| 97 | SR217 | NFS | Lira | 55.2 | 5.1 | 3 |
| 98 | SR252 | LVC | Masaka | 55 | 5.1 | 5 |
| 99 | SR465 | LVC | Sironko | 54.1 | 5 | 5 |
| 100 | SR461 | LVC | Sironko | 53.8 | 1.6 | 3 |
| 101 | SR512 | WNFS | Arua | 53.5 | 1.7 | 5 |
| 102 | SR522 | WMFS | Hoima | 53.2 | 0.4 | 1 |
| 103 | SR33 | NMFS | Kitgum | 53.1 | 2 | 5 |
| 104 | SR432 | WMFS | Hoima | 52.6 | 2.4 | 3 |
| 105 | SR481 | LVC | Sironko | 52.2 | 2 | 2 |
| 106 | SR235 | WNFS | Arua | 52.1 | 4.8 | 5 |
| 107 | SR431 | Unknown | Unknown | 51.3 | 2.8 | 3 |
| 108 | SR468 | LVC | Sironko | 50.4 | 0 | 4 |
| 109 | SR293 | WMFS | Kyenjojo | 50.2 | 4.6 | 3 |
| 110 | SR442 | WMFS | Hoima | 50.1 | 0 | 3 |
| 111 | SR508 | WNFS | Arua | 50 | 2.3 | 4 |
| 112 | SR408 | LVC | Nakaseke | 50 | 1.1 | 2 |
| 113 | SR445 | WMFS | Hoima | 49.6 | 0.4 | 3 |
| 114 | SR462 | LVC | Sironko | 49.5 | 6.5 | 3 |
| 115 | SR67 | LVC | Mukono | 49.3 | 2.8 | 5 |
| 116 | SR3 | TFZ | Amuria | 48.8 | 7 | 1 |
| 117 | SR203 | LVC | Kamuli | 48.3 | 4.5 | 4 |
| 118 | SR249 | Unknown | Unknown | 48.1 | 4.5 | 1 |
| 119 | SR504 | NMFS | Oyam | 47.3 | 0 | 1 |
| 120 | SR290 | WNFS | Kamwenge | 46.9 | 4.3 | 5 |
| 121 | SR283 | WMFS | Kyenjojo | 46.4 | 4.3 | 5 |
| 122 | SR443 | WMFS | Hoima | 45.8 | 0 | 5 |
| 123 | SR302 | WMFS | Hoima | 45.1 | 4.2 | 5 |
| 124 | SR492 | LVC | Sironko | 45.1 | 3.9 | 5 |
| 125 | SR240 | LVC | Rakai | 45 | 4.2 | 3 |
| 126 | SR437 | WMFS | Hoima | 45 | 3 | 4 |
| 127 | SR385 | Unkbown | Unknown | 43.3 | 3 | 3 |
| 128 | SR29 | SWH | Kabale | 42.6 | 3.6 | 1 |
| 129 | SR414 | WMFS | Kabarole | 42.5 | 0 | 2 |
| 130 | SR459 | LVC | Sironko | 42.3 | 3.8 | 3 |
| 131 | SR511 | WNFS | Arua | 41.7 | 5.2 | 5 |
| 132 | SR1 | TFZ | Amuria | 41.4 | 2.1 | 3 |
| 133 | SR336 | NMFS | Oyam | 41.3 | 3.8 | 3 |
| 134 | SR476 | LVC | Sironko | 40.3 | 3.9 | 1 |
| 135 | SR8 | NMFS | Apac | 40.2 | 5 | 5 |
| 136 | SR228 | WNFS | Koboko | 40.1 | 3.7 | 5 |
| 137 | SR41 | LVC | Kayunga | 40.1 | 0 | 5 |
| 138 | SR491 | LVC | Sironko | 39.3 | 3.9 | 3 |
| 139 | SR516 | LVC | Sironko | 39.3 | 1.3 | 3 |
| 140 | SR500 | NMFS | Oyam | 38.5 | 3.4 | 5 |
| 141 | SR63 | LVC | Mukono | 37.1 | 0 | 3 |
| 142 | SR334 | NMFS | Oyam | 36.9 | 3.4 | 3 |
| 143 | SR513 | WNFS | Arua | 36.6 | 1.6 | 5 |
| 144 | SR446 | WMFS | Hoima | 36.4 | 0.9 | 1 |
| 145 | SR495 | LVC | Sironko | 36.3 | 1.3 | 2 |
| 146 | SR476 | LVC | Sironko | 40.3 | 3.9 | 1 |
| 147 | SR8 | NMFS | Apac | 40.2 | 5 | 5 |
| 148 | SR228 | WNFS | Koboko | 40.1 | 3.7 | 5 |
| 149 | SR430 | WMFS | Hoima | 35.9 | 5.2 | 5 |
| 150 | SR411 | WMFS | Hoima | 35.4 | 4.2 | 3 |
| 151 | SR333 | NMFS | Oyam | 35 | 3.2 | 3 |
| 152 | SR521 | WMFS | Hoima | 34.9 | 0 | 3 |
| 153 | SR483 | LVC | Sironko | 33.3 | 0 | 3 |
| 154 | SR321 | NMFS | Oyam | 33.2 | 3.1 | 3 |
| 155 | SR325 | NMFS | Oyam | 33.1 | 3.1 | 3 |
| 156 | SR43 | LVC | Luweero | 32.8 | 0 | 3 |
| 157 | SR485 | LVC | Sironko | 32.3 | 5.1 | 5 |
| 158 | SR209 | LVC | Sironko | 31.9 | 3 | 3 |
| 159 | SR452 | WMFS | Hoima | 31.9 | 0 | 3 |
| 160 | SR433 | WMFS | Hoima | 31.5 | 0 | 1 |
| 161 | SR474 | LVC | Sironko | 30.6 | 1.7 | 1 |
| 162 | SR488 | LVC | Sironko | 29.2 | 0.6 | 3 |
| 163 | SR16 | NMFS | Gulu | 28.6 | 3.5 | 3 |
| 164 | SR460 | LVC | Sironko | 28.5 | 3.5 | 2 |
| 165 | SR4 | TFZ | Amuria | 28.4 | 0.5 | 3 |
| 166 | SR205 | LVC | Jinja | 28.3 | 2.6 | 3 |
| 167 | SR444 | WMFS | Hoima | 27.8 | 2.9 | 1 |
| 168 | SR528 | WMFS | Hoima | 27.4 | 1.5 | 1 |
| 169 | SR69 | LVC | Mubende | 26.1 | 1.7 | 5 |
| 170 | SR200 | EH | Kapchorwa | 24.7 | 2.3 | 1 |
| 171 | SR494 | LVC | Sironko | 24.5 | 3.5 | 2 |
| 172 | SR49 | LVC | Luweero | 24 | 0.9 | 3 |
| 173 | SR484 | LVC | Sironko | 23.2 | 0.5 | 3 |
| 174 | SR256 | Unknown | Unknown | 22.9 | 3.6 | 2 |
| 175 | SR525 | WMFS | Hoima | 22.1 | 0 | 3 |
| 176 | SR515 | LVC | Sironko | 20.9 | 1.3 | 2 |
| 177 | SR448 | WMFS | Hoima | 18.3 | 2.9 | 1 |
| 178 | SR498 | NMFS | Oyam | 18.1 | 0.3 | 3 |
| 179 | SR45 | LVC | Luweero | 18 | 2.9 | 1 |
| 180 | SR493 | LVC | Sironko | 16.8 | 2.1 | 5 |
| 181 | SR435 | WMFS | Hoima | 16.1 | 0 | 4 |
| 182 | SR454 | WMFS | Hoima | 16 | 1.9 | 2 |
| 183 | SR264 | LVC | Bugiri | 15.9 | 1.5 | 3 |
| 184 | SR518 | LVC | Sironko | 15.5 | 1.9 | 2 |
| 185 | SR506 | LVC | Lwengo | 13.2 | 1.5 | 3 |
| 186 | SR9 | NMFS | Apac | 10.2 | 0.1 | 5 |
| 187 | SR489 | LVC | Sironko | 10.1 | 1.4 | 3 |
| 188 | SR475 | LVC | Sironko | 10.1 | 1.1 | 2 |

EH- Eastern Highlands, LVC- Lake Victoria Crescent and Mbale Farmland, NMFS- Northern Mixed Farming System, SWH- South Western Highlands, TFZ- Teso Farming Zone, WMFS- Western Mixed Farming System, WNMFS- West Nile Mixed Fareming System.
