## Supplementary material for "Pathogenic and genetic diversity of *Sclerotium rolfsii,* the causal agent of Southern blight of common bean in Uganda": Table S2.docx

Table S2: Growth rate (cm/day) and number of sclerotia produced by *S. rolfsii* strains. The number of sclerotia was characterized as; 0- none, low- 1 to49, Medium- 50 to 99 and high- 100 and above.

| **S/no** | **Isolate** | **Year** | **District** | **Agroecology** | **Growth**  **Rate** | **SE** | **No. of sclerotia** | **Group** |
| --- | --- | --- | --- | --- | --- | --- | --- | --- |
| 1 | SR468 | 2021 | Sironko | LVC | 3.67 | 0.00 | 290 | High |
| 2 | SR6 | 2013 | Apac | NMFS | 3.63 | 0.01 | 0 | None |
| 3 | SR505 | 2020 | Hoima | WMFS | 3.6 | 0.05 | 99 | Medium |
| 4 | SR495 | 2021 | Sironko | LVC | 3.57 | 0.00 | 0 | None |
| 5 | SR495 | 2021 | Sironko | LVC | 3.57 | 0.07 | 0 | None |
| 6 | SR462 | 2021 | Sironko | LVC | 3.5 | 0.13 | 132 | High |
| 7 | SR462 | 2021 | Sironko | LVC | 3.5 | 0.04 | 132 | High |
| 8 | SR517 | 2021 | Sironko | LVC | 3.47 | 0.01 | 0 | None |
| 9 | SR517 | 2021 | Hoima | WMFS | 3.47 | 0.01 | 0 | Low |
| 10 | SR477 | 2021 | Sironko | LVC | 3.43 | 0.01 | 56 | Medium |
| 11 | SR464 | 2021 | Sironko | LVC | 3.33 | 0.11 | 0 | None |
| 12 | SR446 | 2021 | Hoima | WMFS | 3.33 | 0.04 | 21 | Low |
| 13 | SR446 | 2021 | Hoima | WMFS | 3.33 | 0.01 | 21 | Low |
| 14 | SR205 | 2013 | Jinja | LVC | 3.23 | 0.08 | 171 | Medium |
| 15 | SR472 | 2021 | Sironko | LVC | 3.223 | 0.04 | 0 | None |
| 16 | SR438 | 2021 | Hoima | WMFS | 3.2 | 0.05 | 101 | High |
| 17 | SR438 | 2021 | Hoima | NMFS | 3.2 | 0.06 | 101 | Medium |
| 18 | SR32 | 2013 | Kamuli | LVC | 3.2 | 0.13 | 4 | Low |
| 19 | SR508 | 2021 | Arua | WNFS | 3.18 | 0.03 | 115.7 | medium |
| 20 | SR250 | 2013 | Rakai | LVC | 3.13 | 0.01 | 35 | Low |
| 21 | SR482 | 2021 | Sironko | LVC | 3.13 | 0.03 | 306.3 | High |
| 22 | SR250 | 2013 | Rakai | LVC | 3.13 | 0.00 | 35 | Low |
| 23 | SR450 | 2021 | Hoima | WMFS | 3.1 | 0.00 | 480.7 | High |
| 24 | SR46 | 2013 | Luwero | LVC | 3.1 | 0.03 | 0 | None |
| 25 | SR487 | 2021 | Sironko | LVC | 3.1 | 0.03 | 23 | Low |
| 26 | SR65 | 2013 | Mukono | LVC | 3.1 | 0.11 | 0 | None |
| 27 | SR487 | 2021 | Sironko | LVC | 3.1 | 0.01 | 23 | Low |
| 28 | SR63 | 2013 | Mukono | LVC | 3.1 | 0.05 | 0 | None |
| **S/no** | **Sample** | **Year** | **District** | **Agroecology** | **Growth**  **Rate** | **SE** | **No. of sclerotia** | **Group** |
| 29 | SR53 | 2013 | Mbale | LVC | 3.1 | 0.07 | 0 | None |
| 30 | SR449 | 2021 | Hoima | WMFS | 3.08 | 0.04 | 80 | Medium |
| 31 | SR282 | 2013 | WMFS | Unk | 3.07 | 0.07 | 7.3 | Low |
| 32 | SR203 | 2013 | Kamwenge | WMFS | 3 | 0.04 | 529 | High |
| 33 | SR400 | 2013 | Mbale | LVC | 3 | 0.03 | 0 | None |
| 34 | SR415 | 2013 | Mukono | LVC | 3 | 0.02 | 31.35 | None |
| 35 | SR74 | 2013 | Nakaseke | LVC | 3 | 0.02 | 0 | None |
| 36 | SR491 | 2021 | Sironko | LVC | 3 | 0.06 | 26 | Low |
| 37 | SR45 | 2013 | Luwero | LVC | 3 | 0.08 | 76 | medium |
| 38 | SR74 | 2013 | Nakaseke | LVC | 3 | 0.07 | 0 | None |
| 39 | SR421 | 2013 | Luwero | LVC | 3 | 0.17 | 29 | Low |
| 40 | SR74 | 2013 | Nakaseke | LVC | 3 | 0.03 | 0 | None |
| 41 | SR458 | 2021 | Sironko | LVC | 3 | 0.02 | 60 | Medium |
| 42 | SR475 | 2021 | Sironko | LVC | 2.97 | 0.13 | 47 | Low |
| 43 | SR534 | 2013 | WMFS | LVC | 2.92 | 0.11 | 0 | None |
| 44 | SR534 | 2013 | Mbale | LVC | 2.92 | 0.05 | 0 | None |
| 45 | SR425 | 2013 | Sironko | LVC | 2.91 | 0.01 | 0 | None |
| 46 | SR407 | 2013 | Luwero | LVC | 2.91 | 0.02 | 0 | None |
| 47 | SR67 | 2013 | Mukono | LVC | 2.9 | 0.02 | 0 | None |
| 48 | SR530 | 2021 | Bukoba | Tanzania | 2.9 | 0.05 | 0 | None |
| 49 | SR530 | 2021 | Bukoba | Tanzania | 2.9 | 0.08 | 0 | None |
| 50 | SR290 | 2013 | Kamwenge | WMFS | 2.9 | 0.04 | 54 | Medium |
| 51 | SR484 | 2021 | Sironko | LVC | 2.9 | 0.08 | 217.3 | High |
| 52 | SR139 | 2013 | Sironko | TFZ | 2.9 | 0.06 | 1 | Low |
| 53 | SR408 | 2013 | Nakaseke | LVC | 2.9 | 0.04 | 0 | None |
| 54 | SR52 | 2013 | Luwero | LVC | 2.9 | 0.07 | 0 | None |
| 55 | SR47 | 2013 | Luwero | LVC | 2.9 | 0.04 | 44.35 | Low |
| 56 | SR56 | 2013 | Mbale | LVC | 2.9 | 0.13 | 0 | None |
| 57 | SR252 | 2013 | Masindi | WNFS | 2.88 | 0.00 | 35 | Low |
| 58 | SR501 | 2020 | Oyam | NMFS | 2.87 | 0.00 | 22.3 | Low |
| 59 | SR256 | 2013 | WMFS | WMFS | 2.85 | 0.17 | 9 | Low |
| **S/no** | **Sample** | **Year** | **District** | **Agroecology** | **Growth**  **Rate** | **SE** | **No. of sclerotia** | **Group** |
| 60 | SR494 | 2021 | Sironko | LVC | 2.85 | 0.07 | 0 | None |
| 61 | SR532 | 2013 | Mbale | LVC | 2.85 | 0.04 | 0 | None |
| 62 | SR237 | 2013 | Lwengo | LVC | 2.83 | 0.04 | 256 | High |
| 63 | SR522 | 2021 | Hoima | WMFS | 2.83 | 0.03 | 65 | Medium |
| 64 | SR504 | 2020 | Oyam | NMFS | 2.82 | 0.05 | 248 | High |
| 65 | SR244 | 2013 | WMFS | WMFS | 2.8 | 0.05 | 44 | Low |
| 66 | SR435 | 2021 | Hoima | WMFS | 2.8 | 0.08 | 65 | Medium |
| 67 | SR281 | 2013 | Kabarole | SWH | 2.8 | 0.04 | 476 | High |
| 68 | SR77 | 2013 | Nakaseke | LVC | 2.8 | 0.03 | 0 | None |
| 69 | SR413 | 2013 | Kayunga | LVC | 2.8 | 0.02 | 10.01 | Low |
| 70 | SR45 | 2013 | Luwero | LVC | 2.8 | 0.07 | 76.01 | medium |
| 71 | SR410 | 2013 | Mbale | LVC | 2.8 | 0.05 | 72.01 | medium |
| 72 | SR38 | 2013 | Kyenjojo | WMFS | 2.8 | 0.08 | 0.3 | Low |
| 73 | SR456 | 2021 | Sironko | LVC | 2.78 | 0.02 | 0 | None |
| 74 | SR433 | 2021 | Hoima | WMFS | 2.77 | 0.02 | 64 | Medium |
| 75 | SR283 | 2013 | Kyenjojo | WMFS | 2.77 | 0.04 | 8 | Low |
| 76 | SR249 | 2013 | WMFS | WMFS | 2.76 | 0.04 | 74 | Medium |
| 77 | SR481 | 2021 | Sironko | LVC | 2.75 | 0.01 | 0 | None |
| 78 | SR430 | 2013 | WMFS | WMFS | 2.75 | 0.13 | 4.3 | Low |
| 79 | SR524 | 2021 | Hoima | WMFS | 2.73 | 0.00 | 0 | None |
| 80 | SR518 | 2021 | Hoima | WMFS | 2.73 | 0.04 | 0 | None |
| 81 | SR333 | 2013 | Oyam | NMFS | 2.73 | 0.07 | 18 | Low |
| 82 | SR422 | 2013 | Kayunga | LVC | 2.7 | 0.04 | 0 | None |
| 83 | SR529 | 2021 | Hoima | WMFS | 2.7 | 0.01 | 0 | None |
| 84 | SR302 | 2013 | Hoima | WMFS | 2.7 | 0.04 | 96 | medium |
| 85 | SR525 | 2021 | Hoima | WMFS | 2.67 | 0.06 | 95.3 | Medium |
| 86 | SR55 | 2013 | Mbale | LVC | 2.65 | 0.17 | 0 | None |
| 87 | SR531 | 2021 | Bukoba | Tanzania | 2.63 | 0.07 | 0.7 | Low |
| 88 | SR506 | 2020 | Lwengo | LVC | 2.63 | 0.08 | 0 | None |
| 89 | SR459 | 2021 | Sironko | LVC | 2.6 | 0.08 | 13.3 | low |
| **S/no** | **Sample** | **Year** | **District** | **Agroecology** | **Growth**  **Rate** | **SE** | **No. of sclerotia** | **Group** |
| 90 | SR528 | 2021 | Hoima | WMFS | 2.6 | 0.04 | 43 | low |
| 91 | SR48 | 2013 | Luwero | LVC | 2.6 | 0.04 | 1.01 | low |
| 92 | SR512 | 2021 | Arua | WNFS | 2.6 | 0.01 | 50 | medium |
| 93 | SR454 | 2021 | Hoima | WMFS | 2.57 | 0.04 | 0 | None |
| 94 | SR476 | 2021 | Sironko | LVC | 2.55 | 0.03 | 109.7 | High |
| 95 | SR406 | 2013 | Luwero | LVC | 2.54 | 0.17 | 29.68 | low |
| 96 | SR41 | 2013 | Kyenjojo | WMFS | 2.54 | 0.03 | 0 | None |
| 97 | SR406 | 2013 | Luwero | LVC | 2.53 | 0.01 | 29.68 | Low |
| 98 | SR448 | 2021 | Hoima | WMFS | 2.53 | 0.08 | 0 | None |
| 99 | SR460 | 2021 | Sironko | LVC | 2.52 | 0.00 | 0 | None |
| 100 | SR474 | 2021 | Sironko | LVC | 2.52 | 0.00 | 110 | medium |
| 101 | SR471 | 2021 | Sironko | LVC | 2.5 | 0.02 | 15 | Low |
| 102 | SR70 | 2013 | Mubende | LVC | 2.5 | 0.09 | 332 | High |
| 103 | SR471 | 2021 | Sironko | LVC | 2.5 | 0.01 | 15 | Low |
| 104 | SR439 | 2021 | Hoima | WMFS | 2.5 | 0.13 | 483 | High |
| 105 | SR513 | 2021 | Arua | WNFS | 2.5 | 0.08 | 0 | None |
| 106 | SR488 | 2021 | Sironko | LVC | 2.5 | 0.00 | 1.7 | low |
| 107 | SR35 | 2013 | Kitgum | NMFS | 2.5 | 0.05 | 65 | Medium |
| 108 | SR439 | 2021 | Hoima | WMFS | 2.5 | 0.05 | 483 | High |
| 109 | SR70 | 2013 | Mubende | LVC | 2.5 | 0.01 | 332 | High |
| 110 | SR519 | 2021 | Hoima | WMFS | 2.48 | 0.08 | 40 | Low |
| 111 | SR514 | 2021 | Arua | WNFS | 2.48 | 0.17 | 0 | None |
| 112 | SR502 | 2020 | Oyam | NMFS | 2.48 | 0.02 | 45.3 | Low |
| 113 | SR207 | 2013 | Jinja | LVC | 2.48 | 0.00 | 63 | Medium |
| 114 | SR437 | 2021 | Hoima | WMFS | 2.47 | 0.09 | 276.7 | High |
| 115 | SR5 | 2013 | Amuria | TFZ | 2.45 | 0.07 | 20.68 | low |
| 116 | SR497 | 2020 | Oyam | NMFS | 2.45 | 0.01 | 133.7 | low |
| 117 | SR4 | 2013 | Amuria | TFZ | 2.4 | 0.09 | 0 | None |
| 118 | SR436 | 2021 | Hoima | WMFS | 2.4 | 0.01 | 60 | Medium |
| 119 | SR323 | 2013 | WMFS | WMFS | 2.4 | 0.06 | 37 | Low |
| 120 | SR209 | 2013 | Sironko | LVC | 2.38 | 0.02 | 85 | Medium |
| **S/no** | **Sample** | **Year** | **District** | **Agroecology** | **Growth**  **Rate** | **SE** | **No. of sclerotia** | **Group** |
| 121 | SR287 | 2013 | Mbarara | WMFS | 2.37 | 0.04 | 235 | High |
| 122 | SR485 | 2021 | Sironko | LVC | 2.37 | 0.07 | 0 | None |
| 123 | SR2 | 2013 | Amuria | TFZ | 2.35 | 0.04 | 0 | None |
| 124 | SR515 | 2021 | Hoima | WMFS | 2.35 | 0.02 | 15 | Low |
| 125 | SR356 | 2013 | Lira | NMFS | 2.34 | 0.03 | 112 | medium |
| 126 | SR336 | 2013 | Oyam | NMFS | 2.34 | 0.11 | 32 | Low |
| 127 | SR335 | 2013 | Oyam | NMFS | 2.33 | 0.07 | 2 | low |
| 128 | SR200 | 2013 | Kapchorwa | EH | 2.33 | 0.05 | 311 | High |
| 129 | SR335 | 2013 | Oyam | NMFS | 2.33 | 0.04 | 2 | low |
| 130 | SR332 | 2013 | Oyam | NMFS | 2.3 | 0.05 | 37 | Low |
| 131 | SR8 | 2013 | Apac | NMFS | 2.3 | 0.05 | 0 | None |
| 132 | SR30 | 2013 | Kisoro | SWH | 2.3 | 0.09 | 26.01 | low |
| 133 | SR498 | 2020 | Oyam | NMFS | 2.3 | 0.00 | 119 | medium |
| 134 | SR343 | 2013 | Oyam | NMFS | 2.3 | 0.02 | 2 | Low |
| 135 | SR444 | 2021 | Hoima | WMFS | 2.3 | 0.05 | 90 | Medium |
| 136 | SR321 | 2013 | Oyam | NMFS | 2.3 | 0.17 | 2 | Low |
| 137 | SR334 | 2013 | Oyam | NMFS | 2.3 | 0.13 | 32 | Low |
| 138 | SR466 | 2021 | Sironko | LVC | 2.28 | 0.04 | 0 | medium |
| 139 | SR330 | 2013 | Oyam | NMFS | 2.25 | 0.06 | 58 | Medium |
| 140 | SR141 | 2013 | Wakiso | LVC | 2.25 | 0.01 | 52 | Medium |
| 141 | SR454 | 2021 | Hoima | WMFS | 2.23 | 0.05 | 0 | None |
| 142 | SR45 | 2013 | Luwero | LVC | 2.21 | 0.11 | 76.01 | medium |
| 143 | SR417 | 2013 | Bukedea | TFZ | 2.21 | 0.08 | 0 | None |
| 144 | SR22 | 2013 | Kapchorwa | EH | 2.2 | 0.03 | 366 | High |
| 145 | SR28 | 2013 | Kabale | SWH | 2.2 | 0.01 | 8.35 | low |
| 146 | SR23 | 2013 | Kabale | SWH | 2.2 | 0.08 | 543 | High |
| 147 | SR235 | 2013 | Arua | WNFS | 2.17 | 0.03 | 21 | Low |
| 148 | SR509 | 2021 | Arua | WNFS | 2.15 | 0.00 | 0 | None |
| 149 | SR8 | 2013 | Apac | NMFS | 2.15 | 0.01 | 0 | None |
| 150 | SR527 | 2021 | Hoima | WMFS | 2.12 | 0.11 | 0 | None |
| 151 | SR511 | 2021 | Arua | WNFS | 2.1 | 0.08 | 0 | None |
| **S/no** | **Sample** | **Year** | **District** | **Agroecology** | **Growth**  **Rate** | **SE** | **No. of sclerotia** | **Group** |
| 152 | SR402 | 2013 | Kisoro | LVC | 2.1 | 0.00 | 0 | None |
| 153 | SR25 | 2013 | Kabale | SWH | 2.1 | 0.07 | 0 | None |
| 154 | SR87 | 2021 | Mbale | LVC | 2.1 | 0.00 | 0 | None |
| 155 | SR26 | 2013 | Kabale | SWH | 2.1 | 0.09 | 70.01 | medium |
| 156 | SR492 | 2021 | Sironko | LVC | 2.1 | 0.03 | 0 | None |
| 157 | SR414 | 2013 | Kabarole | SWH | 2.1 | 0.17 | 333 | High |
| 158 | SR29 | 2013 | Kabale | SWH | 2.1 | 0.13 | 60 | medium |
| 159 | SR37 | 2013 | Kyenjojo | WMFS | 2.1 | 0.11 | 49 | Low |
| 160 | SR325 | 2013 | Oyam | NMFS | 2.1 | 0.04 | 2 | Low |
| 161 | SR2 | 2013 | Amuria | TFZ | 2.1 | 0.11 | 0 | None |
| 162 | SR31 | 2013 | Kisoro | SWH | 2.07 | 0.01 | 33.01 | low |
| 163 | SR31 | 2013 | Kisoro | SWH | 2.07 | 0.02 | 33.01 | low |
| 164 | SR31 | 2013 | Kisoro | SWH | 2.07 | 0.01 | 33 | low |
| 165 | SR297 | 2013 | Kabarole | SWH | 2.07 | 0.03 | 248 | High |
| 166 | SR478 | 2021 | Sironko | LVC | 2.05 | 0.01 | 0 | None |
| 167 | SR228 | 2013 | Koboko | WNFS | 2.05 | 0.06 | 58 | Medium |
| 168 | SR510 | 2021 | Arua | WNFS | 2.03 | 0.02 | 0 | None |
| 169 | SR279 | 2013 | Arua | WNFS | 2.03 | 0.07 | 49 | Low |
| 170 | SR443 | 2021 | Hoima | WMFS | 2.03 | 0.07 | 0 | None |
| 171 | SR229 | 2013 | WMFS | WMFS | 2 | 0.08 | 0 | None |
| 172 | SR24 | 2013 | Kabale | SWH | 2 | 0.08 | 33 | low |
| 173 | SR2 | 2013 | Amuria | TFS | 2 | 0.09 | 0 | None |
| 174 | SR4 | 2013 | Amuria | TFZ | 2 | 0.01 | 0 | None |
| 175 | SR201 | 2013 | Kapchorwa | EH | 1.95 | 0.17 | 0 | None |
| 176 | SR523 | 2021 | Hoima | WMFS | 1.9 | 0.06 | 0 | None |
| 177 | SR59 | 2013 | Mbale | LVC | 1.9 | 0.01 | 0 | None |
| 178 | SR11 | 2013 | Apac | NMFS | 1.9 | 0.07 | 0 | None |
| 179 | SR57 | 2013 | Mbale | LVC | 1.9 | 0.00 | 38 | low |
| 180 | SR59 | 2013 | Mbale | LVC | 1.9 | 0.04 | 0 | None |
| 181 | SR225 | 2013 | Arua | WNFS | 1.85 | 0.05 | 17 | Low |
| 182 | SR533 | 2013 | Mbale | LVC | 1.78 | 0.03 | 0 | None |
| **S/no** | **Sample** | **Year** | **District** | **Agroecology** | **Growth**  **Rate** | **SE** | **No. of sclerotia** | **Group** |
| 183 | SR493 | 2021 | Sironko | LVC | 1.73 | 0.07 | 5 | Low |
| 184 | SR493 | 2021 | Sironko | LVC | 1.73 | 0.09 | 5 | Low |
| 185 | SR208 | 2013 | Sironko | LVC | 1.7 | 0.09 | 253 | High |
| 186 | SR33 | 2013 | Kitgum | NMFS | 1.5 | 0.13 | 42 | Low |
| 187 | SR9 | 2013 | Apac | NMFS | 1.25 | 0.08 | 0 | None |
| 188 | SR500 | 2020 | Oyam | NMFS | 1.05 | 0.02 | 0 | None |

EH- Eastern Highlands, LVC- Lake Victoria Crescent and Mbale Farmland, NMFS- Northern Mixed Farming System, SWH- South Western Highlands, TFZ- Teso Farming Zone, WMFS- Western Mixed Farming System, WNMFS- West Nile Mixed Fareming System.
