## Supplementary material for "Pathogenic and genetic diversity of *Sclerotium rolfsii,* the causal agent of Southern blight of common bean in Uganda": Table S3.docx

**Table S3:** Mean and standard errors of morphological and pathogenicity characteristics of *S. rolfsii* strains from five genetically distinct clusters

| Group | Strains | DSI (%) | Growth rate (cm/day) | Average Sclerotia number |
| --- | --- | --- | --- | --- |
| 1 | 35 | 50.4±2.5 b | 2.74±0.05 a | 83 d |
| 2 | 29 | 50.8±2.4 b | 2.66±0.05 a | 36 a |
| 3 | 38 | 49.3±2.4 b | 2..65±0.05 a | 40 a |
| 4 | 32 | 64.9±2.4 a | 2.92±0.05 b | 70 c |
| 5 | 55 | 53.5±2.5 b | 2.79±0.05 c | 28 b |
