## Supplementary material for "Pathogenic and genetic diversity of *Sclerotium rolfsii,* the causal agent of Southern blight of common bean in Uganda": Table S4.docx

Table S4: The proportion of ancestry of *S. rolfsii* isolates and their agroe-cological zones of origin

| S/no | Isolate | Agroecology | Phi cluster 1 | Phi Cluster 2 | Phi cluster 3 | Phi cluster 4 | Phi cluster 5 |
| --- | --- | --- | --- | --- | --- | --- | --- |
| 1 | A01_SR454 | WMFS | 0.6700 | 0.1000 | 0.2300 | 0.0000 | 0.0000 |
| 2 | A02_SR31 | SWH | 0.0000 | 0.9900 | 0.0000 | 0.0100 | 0.0000 |
| 3 | A03_SR67 | LVC | 0.0000 | 0.0000 | 1.0000 | 0.0000 | 0.0000 |
| 4 | A04_SR25 | SWH | 0.8800 | 0.0000 | 0.0000 | 0.0000 | 0.1200 |
| 5 | A05_SR464 | LVC | 0.6400 | 0.1300 | 0.2300 | 0.0000 | 0.0000 |
| 6 | A06_SR400 | LVC | 0.0000 | 0.1700 | 0.0500 | 0.7800 | 0.0000 |
| 7 | A07_SR462 | LVC | 0.0000 | 0.1900 | 0.0200 | 0.0000 | 0.7800 |
| 8 | A08_SR26 | SWH | 0.6600 | 0.1100 | 0.2200 | 0.0000 | 0.0000 |
| 9 | A09_SR482 | LVC | 0.0200 | 0.3000 | 0.0400 | 0.6100 | 0.0300 |
| 10 | A10_SR513 | WNMFS | 0.0000 | 0.0000 | 0.6200 | 0.0000 | 0.3800 |
| 11 | A11_SR422 | LVC | 0.1400 | 0.0000 | 0.0000 | 0.0800 | 0.7800 |
| 12 | A12_SR11 | NMFS | 0.0000 | 0.0000 | 1.0000 | 0.0000 | 0.0000 |
| 13 | B01_SR493 | LVC | 0.0000 | 0.0000 | 1.0000 | 0.0000 | 0.0000 |
| 14 | B02_SR534 | WMFS | 0.0000 | 0.1600 | 0.0400 | 0.7900 | 0.0100 |
| 15 | B03_SR46 | LVC | 0.0400 | 0.0000 | 0.9600 | 0.0000 | 0.0000 |
| 16 | B04_SR475 | LVC | 0.5100 | 0.1300 | 0.3200 | 0.0200 | 0.0200 |
| 17 | B05_SR4 | TFZ | 0.0000 | 0.1900 | 0.0700 | 0.0000 | 0.7400 |
| 18 | B06_SR487 | LVC | 0.0000 | 0.0000 | 0.1000 | 0.0800 | 0.8200 |
| 19 | B07_SR506 | LVC | 0.0000 | 0.0000 | 0.0300 | 0.0300 | 0.9400 |
| 20 | B08_SR472 | LVC | 0.0000 | 0.0000 | 1.0000 | 0.0000 | 0.0000 |
| 21 | B09_SR335 | NMFS | 0.2000 | 0.0000 | 0.0100 | 0.1100 | 0.6800 |
| 22 | B10_SR28 | SWH | 0.0000 | 1.0000 | 0.0000 | 0.0000 | 0.0000 |
| 23 | B11_SR55 | LVC | 0.0000 | 0.0000 | 1.0000 | 0.0000 | 0.0000 |
| 24 | B12_SR534 | WMFS | 0.0000 | 0.1500 | 0.0400 | 0.8000 | 0.0000 |
| 25 | C01_SR33 | NMFS | 0.0300 | 0.0000 | 0.9600 | 0.0000 | 0.0100 |
| 26 | C02_SR70 | NMFS | 0.0000 | 0.0000 | 1.0000 | 0.0000 | 0.0000 |
| 27 | C03_SR476 | LVC | 0.0200 | 0.2900 | 0.0400 | 0.6100 | 0.0300 |
| 28 | C04_SR530 | TANZANIA | 0.8600 | 0.0000 | 0.0000 | 0.0300 | 0.1100 |
| 29 | C05_SR2 | TFZ | 0.6700 | 0.0900 | 0.2400 | 0.0000 | 0.0000 |
| 30 | C06_SR494 | LVC | 0.9300 | 0.0000 | 0.0000 | 0.0000 | 0.0700 |
| 31 | C07_SR141 | LVC | 0.0700 | 0.0000 | 0.9300 | 0.0000 | 0.0000 |
| 32 | C08_SR514 | WNMFS | 0.0000 | 0.0000 | 1.0000 | 0.0000 | 0.0000 |
| 33 | C09_SR504 | NMFS | 0.0000 | 0.1700 | 0.0600 | 0.7600 | 0.0100 |
| 34 | C10_SR74 | LVC | 0.0500 | 0.0100 | 0.9300 | 0.0000 | 0.0100 |
| 35 | C11_SR491 | LVC | 0.0000 | 0.2000 | 0.0500 | 0.0000 | 0.7500 |
| 36 | C12_SR448 | WMFS | 0.1300 | 0.0000 | 0.0000 | 0.8700 | 0.0000 |
| 37 | D01_SR511 | WNMFS | 0.0000 | 0.0000 | 0.6200 | 0.0000 | 0.3800 |
| 38 | D02_SR449 | WMFS | 0.9400 | 0.0000 | 0.0000 | 0.0000 | 0.0600 |
| 39 | D03_SR531 | TANZANIA | 0.8500 | 0.0000 | 0.0000 | 0.0300 | 0.1100 |
| 40 | D04_SR406 | LVC | 0.0000 | 0.1700 | 0.0500 | 0.7900 | 0.0000 |
| 41 | D05_SR256 | WMFS | 0.6600 | 0.0900 | 0.2500 | 0.0000 | 0.0000 |
| 42 | D06_SR332 | NMFS | 0.0000 | 0.0000 | 1.0000 | 0.0000 | 0.0000 |
| 43 | D07_SR500 | NMFS | 0.0000 | 0.0000 | 1.0000 | 0.0000 | 0.0000 |
| 44 | D08_SR525 | WMFS | 0.1200 | 0.0000 | 0.0000 | 0.0700 | 0.8200 |
| 45 | D09_SR435 | WMFS | 0.0200 | 0.9700 | 0.0000 | 0.0100 | 0.0000 |
| 46 | D10_SR488 | LVC | 0.0000 | 0.0000 | 0.0400 | 0.0400 | 0.9200 |
| 47 | D11_SR529 | WMFS | 0.0000 | 0.0000 | 1.0000 | 0.0000 | 0.0000 |
| 48 | D12_SR518 | LVC | 0.8900 | 0.0000 | 0.0000 | 0.0000 | 0.1000 |
| 49 | E01_SR250 | LVC | 0.0000 | 0.0000 | 0.0000 | 0.0300 | 0.9700 |
| 50 | E02_SR406 | LVC | 0.0000 | 0.1700 | 0.0500 | 0.7800 | 0.0000 |
| 51 | E03_SR530 | TANZANIA | 0.8500 | 0.0000 | 0.0000 | 0.0300 | 0.1200 |
| 52 | E04_SR510 | WNMFS | 0.6400 | 0.1100 | 0.2500 | 0.0000 | 0.0000 |
| 53 | E05_SR330 | NMFS | 0.0000 | 1.0000 | 0.0000 | 0.0000 | 0.0000 |
| 54 | E06_SR229 | LVC | 0.0200 | 0.9600 | 0.0000 | 0.0200 | 0.0000 |
| 55 | E07_SR509 | WNMFS | 0.0000 | 0.0000 | 0.6200 | 0.0000 | 0.3800 |
| 56 | E08_SR481 | LVC | 0.6500 | 0.0900 | 0.2600 | 0.0000 | 0.0000 |
| 57 | E09_SR532 | WMFS | 0.0000 | 0.1500 | 0.0500 | 0.7800 | 0.0200 |
| 58 | E10_SR524 | WMFS | 0.0900 | 0.0000 | 0.0000 | 0.0600 | 0.8500 |
| 59 | E11_SR35 | NMFS | 0.0000 | 0.1700 | 0.0000 | 0.0000 | 0.8300 |
| 60 | E12_SR281 | WNMFS | 0.0400 | 0.0000 | 0.9600 | 0.0000 | 0.0000 |
| 61 | F01_SR471 | LVC | 0.0000 | 0.0000 | 0.0700 | 0.0600 | 0.8700 |
| 62 | F02_SR523 | WMFS | 0.0000 | 0.0000 | 1.0000 | 0.0000 | 0.0000 |
| 63 | F03_SR519 | WMFS | 0.0000 | 1.0000 | 0.0000 | 0.0000 | 0.0000 |
| 64 | F04_SR495 | LVC | 0.6400 | 0.1100 | 0.2500 | 0.0000 | 0.0000 |
| 65 | F05_SR517 | LVC | 0.8900 | 0.0000 | 0.0000 | 0.0000 | 0.1100 |
| 66 | F06_SR287 | WMFS | 0.0000 | 0.1500 | 0.0400 | 0.8000 | 0.0100 |
| 67 | F07_SR87 | NMFS | 0.0200 | 0.0100 | 0.9600 | 0.0000 | 0.0100 |
| 68 | F08_SR8 | NMFS | 0.0300 | 0.0000 | 0.9500 | 0.0000 | 0.0100 |
| 69 | F09_SR446 | WMFS | 0.1500 | 0.0000 | 0.0000 | 0.8500 | 0.0000 |
| 70 | F10_SR517 | LVC | 0.9000 | 0.0000 | 0.0000 | 0.0000 | 0.1000 |
| 71 | F11_SR502 | NMFS | 0.0600 | 0.0100 | 0.9300 | 0.0000 | 0.0000 |
| 72 | F12_SR279 | WNMFS | 0.0500 | 0.0000 | 0.9400 | 0.0000 | 0.0000 |
| 73 | G01_SR402 | SWH | 0.0000 | 1.0000 | 0.0000 | 0.0000 | 0.0000 |
| 74 | G02_SR59 | LVC | 0.9000 | 0.0000 | 0.0000 | 0.0000 | 0.1000 |
| 75 | G03_SR249 | NMFS | 0.1000 | 0.0000 | 0.0000 | 0.8900 | 0.0000 |
| 76 | G04_SR460 | LVC | 0.6600 | 0.1000 | 0.2400 | 0.0000 | 0.0000 |
| 77 | G05_SR244 | NMFS | 0.0100 | 0.9900 | 0.0000 | 0.0000 | 0.0000 |
| 78 | G06_SR290 | WMFS | 0.0600 | 0.0000 | 0.9400 | 0.0000 | 0.0000 |
| 79 | G07_SR425 | LVC | 0.0000 | 0.1700 | 0.0000 | 0.0000 | 0.8300 |
| 80 | G08_SR415 | LVC | 0.0500 | 0.0100 | 0.9300 | 0.0000 | 0.0100 |
| 81 | G09_SR282 | WNMFS | 0.0100 | 0.9800 | 0.0000 | 0.0100 | 0.0000 |
| 82 | G10_SR65 | LVC | 0.0500 | 0.0100 | 0.9400 | 0.0000 | 0.0000 |
| 83 | G11_SR492 | LVC | 0.0000 | 0.0000 | 1.0000 | 0.0000 | 0.0000 |
| 84 | G12_SR430 | WMFS | 0.0200 | 0.0000 | 0.9800 | 0.0000 | 0.0000 |
| 85 | H01_SR450 | WMFS | 0.0000 | 0.2100 | 0.0400 | 0.7500 | 0.0000 |
| 86 | H02_SR438 | WMFS | 0.0300 | 0.9600 | 0.0000 | 0.0000 | 0.0100 |
| 87 | H03_SR203 | LVC | 0.0000 | 1.0000 | 0.0000 | 0.0000 | 0.0000 |
| 88 | H04_SR471 | LVC | 0.6800 | 0.1000 | 0.2300 | 0.0000 | 0.0000 |
| 89 | H05_SR515 | LVC | 0.8900 | 0.0000 | 0.0000 | 0.0000 | 0.1100 |
| 90 | H06_SR5 | TFZ | 0.0200 | 0.0000 | 0.9800 | 0.0000 | 0.0000 |
| 91 | H07_SR45 | LVC | 0.0000 | 0.1700 | 0.0600 | 0.7700 | 0.0000 |
| 92 | H08_SR22 | NMFS | 0.1100 | 0.0000 | 0.0000 | 0.8900 | 0.0000 |
| 93 | H09_SR439 | WMFS | 0.0000 | 1.0000 | 0.0000 | 0.0000 | 0.0000 |
| 94 | H10_SR30 | WMFS | 0.0000 | 1.0000 | 0.0000 | 0.0000 | 0.0000 |
| 95 | H11_SR41 | LVC | 0.0400 | 0.0100 | 0.9400 | 0.0000 | 0.0100 |
| 96 | H12_SR459 | LVC | 0.1100 | 0.0000 | 0.0000 | 0.0600 | 0.8400 |
| 97 | A01_SR436 | WMFS | 0.0500 | 0.8900 | 0.0100 | 0.0400 | 0.0100 |
| 98 | A02_SR201 | EH | 0.0000 | 0.1500 | 0.0500 | 0.7900 | 0.0100 |
| 99 | A03_SR454 | WMFS | 0.0000 | 0.0000 | 1.0000 | 0.0000 | 0.0000 |
| 100 | A04_SR343 | LVC | 0.0000 | 1.0000 | 0.0000 | 0.0000 | 0.0000 |
| 101 | A05_SR139 | WNMFS | 0.0000 | 0.9900 | 0.0000 | 0.0100 | 0.0000 |
| 102 | A06_SR45 | LVC | 0.1100 | 0.0000 | 0.0000 | 0.8900 | 0.0000 |
| 103 | A07_SR57 | LVC | 0.0000 | 0.1700 | 0.0000 | 0.0000 | 0.8300 |
| 104 | A08_SR477 | LVC | 0.0000 | 0.1500 | 0.0500 | 0.7800 | 0.0100 |
| 105 | A09_SR48 | LVC | 0.9400 | 0.0000 | 0.0000 | 0.0000 | 0.0600 |
| 106 | A10_SR252 | LVC | 0.0000 | 0.0000 | 1.0000 | 0.0000 | 0.0000 |
| 107 | A11_SR200 | EH | 0.0000 | 0.1600 | 0.0700 | 0.7400 | 0.0300 |
| 108 | A12_SR335 | NMFS | 0.1500 | 0.0000 | 0.0000 | 0.0800 | 0.7700 |
| 109 | B01_SR433 | WMFS | 0.1000 | 0.0000 | 0.0000 | 0.9000 | 0.0000 |
| 110 | B02_SR438 | WMFS | 0.0000 | 1.0000 | 0.0000 | 0.0000 | 0.0000 |
| 111 | B03_SR417 | TFZ | 0.0000 | 0.1600 | 0.0000 | 0.0000 | 0.8400 |
| 112 | B04_SR474 | LVC | 0.0100 | 0.3000 | 0.0400 | 0.6200 | 0.0300 |
| 113 | B05_SR70 | LVC | 0.0000 | 0.0000 | 1.0000 | 0.0000 | 0.0000 |
| 114 | B06_SR462 | LVC | 0.0000 | 0.1900 | 0.0200 | 0.0000 | 0.7900 |
| 115 | B07_SR468 | LVC | 0.0000 | 1.0000 | 0.0000 | 0.0000 | 0.0000 |
| 116 | B08_SR444 | WMFS | 0.1300 | 0.0000 | 0.0000 | 0.8700 | 0.0000 |
| 117 | B09_SR47 | LVC | 0.0000 | 0.1600 | 0.0600 | 0.7700 | 0.0100 |
| 118 | B10_SR512 | WNMFS | 0.0000 | 0.0000 | 0.6200 | 0.0000 | 0.3800 |
| 119 | B11_SR458 | LVC | 0.0000 | 0.0000 | 0.0100 | 0.0400 | 0.9500 |
| 120 | B12_SR495 | LVC | 0.0100 | 0.3100 | 0.0400 | 0.6100 | 0.0300 |
| 121 | C01_SR250 | WMFS | 0.0000 | 0.0000 | 1.0000 | 0.0000 | 0.0000 |
| 122 | C02_SR446 | WMFS | 0.1100 | 0.0000 | 0.0000 | 0.8900 | 0.0000 |
| 123 | C03_SR466 | LVC | 0.0000 | 0.1800 | 0.0000 | 0.0000 | 0.8200 |
| 124 | C04_SR501 | NMFS | 0.1100 | 0.0000 | 0.0000 | 0.8900 | 0.0000 |
| 125 | C05_SR63 | LVC | 0.0000 | 0.0000 | 0.0100 | 0.0300 | 0.9500 |
| 126 | C06_SR408 | LVC | 0.9200 | 0.0000 | 0.0000 | 0.0000 | 0.0800 |
| 127 | C07_SR8 | NMFS | 0.0200 | 0.0000 | 0.9600 | 0.0000 | 0.0100 |
| 128 | C08_SR413 | LVC | 0.0400 | 0.0000 | 0.9600 | 0.0000 | 0.0000 |
| 129 | C09_SR45 | LVC | 0.6500 | 0.1000 | 0.2500 | 0.0000 | 0.0000 |
| 130 | C10_SR2 | TFZ | 0.0000 | 0.0000 | 1.0000 | 0.0000 | 0.0000 |
| 131 | C11_SR522 | WMFS | 0.0000 | 0.1600 | 0.0500 | 0.7700 | 0.0200 |
| 132 | C12_SR334 | NMFS | 0.1600 | 0.0000 | 0.0000 | 0.0800 | 0.7600 |
| 133 | D01_SR498 | NMFS | 0.0000 | 0.0000 | 0.0300 | 0.0500 | 0.9200 |
| 134 | D02_SR439 | WMFS | 0.0600 | 0.8900 | 0.0100 | 0.0300 | 0.0100 |
| 135 | D03_SR302 | WMFS | 0.0000 | 0.0000 | 1.0000 | 0.0000 | 0.0000 |
| 136 | D04_SR487 | LVC | 0.0000 | 0.0000 | 0.0600 | 0.0800 | 0.8600 |
| 137 | D05_SR456 | LVC | 0.0000 | 0.0000 | 1.0000 | 0.0000 | 0.0000 |
| 138 | D06_SR74 | LVC | 0.0100 | 0.9800 | 0.0000 | 0.0000 | 0.0000 |
| 139 | D07_SR37 | WMFS | 0.0000 | 1.0000 | 0.0000 | 0.0000 | 0.0000 |
| 140 | D08_SR235 | WNMFS | 0.0000 | 0.0000 | 0.6200 | 0.0000 | 0.3800 |
| 141 | D09_SR56 | LVC | 0.6600 | 0.1200 | 0.2300 | 0.0000 | 0.0000 |
| 142 | D10_SR208 | LVC | 0.0600 | 0.8900 | 0.0100 | 0.0300 | 0.0100 |
| 143 | D11_SR297 | WMFS | 0.0100 | 0.9700 | 0.0000 | 0.0100 | 0.0000 |
| 144 | D12_SR205 | LVC | 0.0000 | 0.1700 | 0.0000 | 0.0000 | 0.8300 |
| 145 | E01_SR478 | LVC | 0.0200 | 0.0100 | 0.9600 | 0.0000 | 0.0100 |
| 146 | E02_SR31 | SWH | 0.0000 | 1.0000 | 0.0000 | 0.0000 | 0.0000 |
| 147 | E03_SR443 | WMFS | 0.0000 | 0.0000 | 1.0000 | 0.0000 | 0.0000 |
| 148 | E04_SR336 | NMFS | 0.1300 | 0.0000 | 0.0000 | 0.0700 | 0.8000 |
| 149 | E05_SR77 | LVC | 0.4100 | 0.0000 | 0.1900 | 0.1500 | 0.2500 |
| 150 | E06_SR29 | SWH | 0.1000 | 0.0000 | 0.0000 | 0.9000 | 0.0000 |
| 151 | E07_SR2 | TFZ | 0.0000 | 0.1600 | 0.0000 | 0.0000 | 0.8400 |
| 152 | E08_SR74 | LVC | 0.0400 | 0.0100 | 0.9500 | 0.0000 | 0.0000 |
| 153 | E09_SR9 | NMFS | 0.0000 | 0.0000 | 1.0000 | 0.0000 | 0.0000 |
| 154 | E10_SR59 | LVC | 0.8900 | 0.0000 | 0.0000 | 0.0000 | 0.1100 |
| 155 | E11_SR53 | NMFS | 0.1100 | 0.0000 | 0.0000 | 0.8900 | 0.0000 |
| 156 | E12_SR281_P1 | WNMFS | 0.0200 | 0.0000 | 0.9800 | 0.0000 | 0.0000 |
| 157 | F01_SR527 | WMFS | 0.0200 | 0.0000 | 0.9700 | 0.0000 | 0.0000 |
| 158 | F02_SR508 | WNMFS | 0.0100 | 0.9900 | 0.0000 | 0.0000 | 0.0100 |
| 159 | F03_SR32 | LVC | 0.0000 | 0.0000 | 1.0000 | 0.0000 | 0.0000 |
| 160 | F04_SR437 | WMFS | 0.0000 | 1.0000 | 0.0000 | 0.0000 | 0.0000 |
| 161 | F05_SR533 | WMFS | 0.0000 | 0.0000 | 0.0200 | 0.0400 | 0.9300 |
| 162 | F06_SR24 | SWH | 0.0000 | 1.0000 | 0.0000 | 0.0000 | 0.0000 |
| 163 | F07_SR325 | NMFS | 0.0000 | 0.0000 | 0.0100 | 0.0300 | 0.9600 |
| 164 | F08_SR52 | LVC | 0.0000 | 0.0000 | 0.1200 | 0.0800 | 0.8000 |
| 165 | F09_SR4 | TFZ | 0.0000 | 0.1700 | 0.0100 | 0.0000 | 0.8200 |
| 166 | F10_SR321 | NMFS | 0.0000 | 0.1700 | 0.0000 | 0.0000 | 0.8300 |
| 167 | F11_SR410 | LVC | 0.0000 | 0.1600 | 0.0600 | 0.7700 | 0.0100 |
| 168 | G01_SR493 | LVC | 0.0000 | 0.0000 | 1.0000 | 0.0000 | 0.0000 |
| 169 | G02_SR356 | NMFS | 0.0000 | 1.0000 | 0.0000 | 0.0000 | 0.0000 |
| 170 | G03_SR484 | LVC | 0.0000 | 0.1800 | 0.0000 | 0.0000 | 0.8100 |
| 171 | G04_SR528 | WMFS | 0.1000 | 0.0000 | 0.0000 | 0.9000 | 0.0000 |
| 172 | G05_SR485 | LVC | 0.0200 | 0.0000 | 0.9800 | 0.0000 | 0.0000 |
| 173 | G06_SR31 | SWH | 0.0000 | 1.0000 | 0.0000 | 0.0000 | 0.0000 |
| 174 | G07_SR421 | LVC | 0.0000 | 0.1900 | 0.0000 | 0.0000 | 0.8100 |
| 175 | G08_SR225 | WNMFS | 0.0100 | 0.9900 | 0.0000 | 0.0000 | 0.0000 |
| 176 | G09_SR407 | LVC | 0.1200 | 0.0000 | 0.0000 | 0.0700 | 0.8100 |
| 177 | G10_SR228 | WNMFS | 0.0500 | 0.0000 | 0.9500 | 0.0000 | 0.0000 |
| 178 | G11_SR38 | WMFS | 0.0600 | 0.0000 | 0.9300 | 0.0000 | 0.0000 |
| 179 | H01_SR237 | LVC | 0.0000 | 1.0000 | 0.0000 | 0.0000 | 0.0000 |
| 180 | H02_SR333 | NMFS | 0.1100 | 0.0000 | 0.0000 | 0.0600 | 0.8300 |
| 181 | H03_SR497 | WMFS | 0.0000 | 0.1700 | 0.0500 | 0.7800 | 0.0000 |
| 182 | H04_SR414 | WMFS | 0.9000 | 0.0000 | 0.0000 | 0.0000 | 0.1000 |
| 183 | H05_SR505 | LVC | 0.0000 | 0.1600 | 0.0500 | 0.7800 | 0.0000 |
| 184 | H06_SR209 | LVC | 0.1200 | 0.0000 | 0.0000 | 0.0600 | 0.8200 |
| 185 | H07_SR323 | LVC | 0.0000 | 0.0000 | 1.0000 | 0.0000 | 0.0000 |
| 186 | H08_SR23 | SWH | 0.0000 | 1.0000 | 0.0000 | 0.0000 | 0.0000 |
| 187 | H09_SR207 | LVC | 0.0000 | 0.0000 | 1.0000 | 0.0000 | 0.0000 |
| 188 | H10_SR6 | NMFS | 0.0000 | 0.1700 | 0.0500 | 0.7800 | 0.0000 |
| 189 | H11_SR283 | WMFS | 0.0000 | 0.0000 | 1.0000 | 0.0000 | 0.0000 |

EH- Eastern Highlands, LVC- Lake Victoria Crescent and Mbale Farmland, NMFS- Northern Mixed Farming System, SWH- South Western Highlands, TFZ- Teso Farming Zone, WMFS- Western Mixed Farming System, WNMFS- West Nile Mixed Fareming System.
